# Longitudinal assessment of water-reaching reveals altered cortical activity and fine motor coordination defects in a Huntington Disease model

**DOI:** 10.1101/2022.09.02.505959

**Authors:** Yundi Wang, Marja D. Sepers, Dongsheng Xiao, Lynn A. Raymond, Timothy H. Murphy

## Abstract

Huntington Disease (HD), caused by dominantly inherited expansions of a CAG repeat results in characteristic motor dysfunction. Although gross motor and balance defects have been extensively characterized in multiple HD mouse models using tasks such as rotarod, beam walking and gait analysis, little is known about forelimb deficits. Here we use a high-throughput alternating reward/non-reward water-reaching task conducted daily over ∼2 months to simultaneously monitor forelimb impairment and mesoscale cortical changes in GCaMP activity, comparing female zQ175 (HD) and wildtype (WT) littermate mice, starting at ∼5.5 months of age. Behavioral analysis of the water-reaching task reveals that HD mice, despite learning the water-reaching task as proficiently as WT mice, take longer to learn the alternating event sequence. Although WT mice displayed no significant changes in cortical activity and reaching trajectory throughout the testing period, HD mice exhibited an increase in cortical activity – especially in the secondary motor and retrosplenial cortices – over time, as well as longer and more variable reaching trajectories by ∼7 months of age. HD mice also experienced a progressive reduction in successful performance rates. Tapered beam and rotarod tests before and/or after water-reaching assessment confirmed these early and manifest stages of HD characterized by the absence and presence of failed water-reaching trials, respectively. Reduced DARPP-32 (marker for striatal medium spiny neurons) expression in HD mice further confirmed disease pathology. The water-reaching task can be used to inform HD and potentially other movement disorder onset, therapeutic intervention windows and test drug efficacy.

**Significance statement:** The movement disorder, Huntington Disease (HD), has been extensively studied in preclinical settings using mouse models of disease examining gross motor and balance defects. Little however, is known regarding forelimb deficits and underlying cortical circuit changes. Using a high-throughput alternating reward/non-reward water-reaching task, we characterized early event sequence learning defects in HD mice aged ∼5.5 months. Progressive forelimb movement defects first become apparent at ∼6.5 months of age with corresponding increases in cortical activity associated with reaching observed over time. These forelimb defects revealed in the water-reaching task are coincident with gross motor defects characterized using the tapered beam and rotarod tasks, demonstrating the suitability of the water-reaching task in phenotyping HD motor deficits.

## Introduction

Synaptic and circuit changes that precede progressive striatal medium spiny neuron (MSN) and cortical neuronal loss, in the case of Huntington Disease (HD) results in a characteristic triad of symptoms: motor dysfunction, cognitive impairment, and neuropsychiatric symptoms (McColgan and Tabrizi, 2018; Cepeda and Levine, 2022). These hallmark motor symptoms of HD include fine motor incoordination, chorea, bradykinesia, rigidity and difficulties with balance and gait.

Since the discovery that dominantly inherited expansions >39 of a CAG triplet repeat in exon 1 of the *HTT* gene causes HD (MacDonald et al., 1993), over 50 distinct mouse and rat models with increasingly better face and construct validity have been developed (Pouladi et al., 2013; Menalled et al., 2014). Although several assessments such as rotarod, balance beam and gait tasks (such as the footprint test) are commonly used to assess motor defects in HD mice (Pallier et al., 2009; Brooks et al., 2012; Abada et al., 2013; Southwell et al., 2016), bodyweight remains a major confound of these tasks (McFadyen et al., 2003; Batka et al., 2014) necessitating the development and usage of other behavioral assessments.

In clinical settings, impairments in reaching and/or skilled hand movements have been observed in HD patients (Klein et al., 2011). Skilled forelimb movement learning and performance also has been examined in HD mice using automated home-cage lever pulling systems (Woodard et al., 2017, 2021), demonstrating that HD motor learning deficits are related to impaired striatal neuronal plasticity (Woodard et al., 2021). Given reaching towards a target and manipulating objects is commonly used in our daily lives and the general ‘reach-to-grasp’ features of forelimb movements has been shown to be similar between humans and rodents (Galiñanes et al., 2018), preclinical behavioral assessment of skilled forelimb reaching tasks could improve our understanding of HD movement defects. Water-reaching tasks further enable longitudinal multi-trial assessment providing a high throughput system for behavioral phenotyping throughout the duration of disease progression, and may detect more subtle motor learning and movement defects. Combining water-reaching assessment with simultaneous recording of widefield cortical activity further enables examination of pathophysiological circuit changes underlying HD movement defects.

To date, few studies exist examining these wide-scale cortical circuit changes in HD mouse models. Using 3D magnetic resonance imaging, arteriolar cerebral blood volume level changes in the striatum and motor cortex were observed in HD mice beginning at 3 months of age which worsened overtime (Liu et al., 2021). Hemodynamic measurements are, however, indirect indicators of neuronal activity. Using mesoscale voltage-sensitive dye imaging, our group has shown that hindlimb stimulation evokes a larger area and longer lasting cortical response in anesthetized HD compared to WT mice (Sepers et al., 2021). Given recent neurophysiology studies have demonstrated the involvement of multiple brain regions in sensation, cognition and movement (Pinto et al., 2019; Steinmetz et al., 2019), widefield functional assessment of neuronal circuit changes during task performance is needed throughout the time course of HD disease pathology.

In this study, we used a water-reaching task to demonstrate progressive changes in widefield cortical activity and skilled forelimb movement defects. Motor deficits in the water-reaching task correlated with stage-dependent deficits on tapered beam and rotarod tests as well as post-hoc immunohistochemistry staining for striatal MSNs. The full-length *HTT* knock-in heterozygous zQ175 mouse model was employed due to its advantage of having a relatively slow disease development and greater construct validity over other HD mouse models (e.g. R6/1, R6/2 and BACHD)(Pouladi et al., 2013).

## Methods

### Animals

All experiments and procedures were carried out in accordance with the Canadian Council on Animal Care and approved by the University of British Columbia Committee on Animal Care (protocols A18-0036 and A19-0076). Mice were group housed with 2 to 4 mice per cage under a controlled 12 hr light/dark cycle (7:00 lights on, 19:00 lights off). Standard laboratory mouse diet was available *ad libitum*. Water was available *ad libitum* except during the duration of head-fixed water-reaching behavioral testing and when mice were readjusted to *ad libitum* water consumption. Surgery and subsequent behavioral testing was performed on 6 female heterozygous zQ175 knock-in C57BL/6 mice expressing GCaMP6s and 6 female wildtype (WT) littermates as controls starting at ∼5 months of age. zQ175 C57BL/6 mice and WT littermates, both expressing GCaMP6s, were obtained by first crossing heterozygous zQ175 C57BL/6 mice (https://www.jax.org/strain/029928) with homozygous transgenic Thy-1 GCaMP6s line 4.3 C57BL/6 mice (HHMI Janelia Research Campus)(Dana et al., 2014; Sofroniew et al., 2016) then crossing subsequent offspring to obtain homozygous GCaMP6s expression. Animal tissue was collected through ear clipping at weaning. DNA extraction and PCR analysis were subsequently used to determine genotype. Health status and weight of all animals was assessed daily.

### Animal surgery

All mice were subjected to head-bar and chronic transcranial window surgery as previously described (Murphy et al., 2016; Silasi et al., 2016) and allowed to recover for a week before commencement of behavioral testing. Briefly, an incision and skin retraction over the cortex was made enabling a glass coverslip to be applied using Metabond clear dental cement (Parkell, Edgewood, NY, USA; Product: C&B Metabond) onto un-thinned bone. A steel head-fixation bar was also placed 4 mm posterior between bregma and the bar edge.

### Behavioral testing timeline

During the handling period, mice were first habituated to daily human contact for three weeks. Mice then underwent surgery and were allowed to recover for one week before initial tapered beam assessment (5 days). After 2 days of initial tapered beam testing, mice were also habituated to first the confinement tube only, then to the confinement tube and head fixation and, finally the confinement tube, head fixation and experimental setup. The duration of head fixation was progressively increased at a rate of ∼7 min/day for 5 days.

Mice were then water restricted for skilled forelimb head-fixed water-reaching behavioral training and testing. From the water-reaching task, mice had the potential to receive ∼1 mL/day of water. Given variation in weight due to *ad libitum* food consumption and excrement, mice who either performed poorly or lost more than 0.5 g in weight compared to the previous day were given up to ∼1.1 mL of task-independent water. All mice received ∼100 µL of additional water daily. As such, all mice received ∼1.1 µL of task-independent and/or behavioral test-derived water daily. The humane endpoint was defined as a maximal weight loss of 15% from a pre-water-restricted baseline weight. No mice reached the humane endpoint during the duration of the study.

After a maximum of 67 days of water restriction and water-reaching behavioral training and testing, mice were readjusted to *ad libitum* access to water. During this readjustment period, mice received 1.1 mL of task-independent water on the first day. In subsequent days mice received progressively increased water at a rate of ∼500 μL/day for ∼4 days. This additional water was administered at three different times during the light cycle to prevent water intoxication. After stabilization of mouse weight, *ad libitum* access to water was restored for the duration of the behavioral testing.

Accelerating rotarod (4 days) and final tapered beam testing (7 days) was then conducted with a 1 day recovery period between the two behavior assessments. All animals were sacrificed with intraperitoneal injection of pentobarbital sodium (240 mg/kg) and transcardially perfused with first 10 mL phosphate-buffered saline (PBS) then 10 mL 10% neutral buffered formalin (NBF). Whole brains were removed and post-fixed in NBF for post-mortem immunohistochemistry.

### Head-fixed water-reaching test

Mice underwent head-bar and chronic transcranial surgery and were trained to reach for water under head-fixed conditions following in part a previously described protocol (Galiñanes et al., 2018). All mice were water restricted after habituation to the confinement tube, head-fixation and experimental set up. A platform which extends 1.5 cm from the base of the confinement tube allowed the mice to rest their forepaws while not reaching for water. The water spout was fashioned using a blunted 22G needle bent at a 90° angle. The starting position of the water spout was ∼0.75-1 cm posterior from the tip of the snout and positioned laterally so that the water drop made minimal contact with the whiskers. At this position, mice could touch the water spout with their paws and feel the water drop if they groomed which transitioned to reaching. For mice which did not groom, the water drop was allowed to touch their whisker pad which promoted grooming and transition into reaching toward the spout. Once mice started reaching, the distance of the water spout was gradually increased until a final distance of ∼1.5 cm lateral and ∼0.5 cm posterior to the tip of the snout was achieved. Only mice that reached the final distance with a success rate of at least ∼80% on Day 15 were included for further analysis.

Trial structure included alternating reward and no reward trials. Unless an electronic failure occurred, all trials started with a rewarded trial. The experimental setup was illuminated with infrared LED illuminator lights. A Raspberry Pi single-board computer and custom Python script was used to control camera recording, blue and green light used to illuminate the cortex, cue light signal, cue buzzer signal, water solenoid to deliver the water reward and capacitive touch sensor connected to the water spout.

At the start of reward and no reward trials, a Raspberry Pi infrared night vision camera (320 x 320 pixels; 60 Hz) and a 1M60 Pantera CCD camera (Dalsa) enabled behavioral and GCaMP activity recording, respectively. The cortex was illuminated using alternating green and blue light providing information about hemodynamic changes and exciting GCaMP, respectively and collected as 12-bit images through the Dalsa camera using XCAP imaging software (120 Hz). Binning camera pixels (8 x 8) produced a resolution of 68 µm/pixel. These imaging parameters have been used previously for widefield cortical GCaMP imaging (Vanni and Murphy, 2014; Xiao et al., 2017).

A 0.1 s duration green LED light flash 2 s after the start of camera recording was followed by a 0.1 s buzzer tone in the case of non-rewarded trials or a buzzer tone combined with simultaneous ∼20 µL water reward in the case of rewarded trials 6 s after the start of camera recording. For rewarded trials, if a spout touch was detected by the capacitive touch sensor (Adafruit Industries, New York, NY, USA) after delivery of the water reward, the Picamera and Dalsa camera recording would cease 4 s after the time of spout touch. If a touch was not detected or it was a non-rewarded trial, camera recording would cease 10 s after delivery of the reward. Rewarded trials therefore ranged from 10-16 s and non-rewarded trials were 16 s in length. The intertrial interval was 4 s. A total of ∼120 trials over a duration of ∼39 min were conducted daily for a maximum of ∼67 days.

Since the capacitive touch sensor was found to not accurately determine reaches, touches and/or contact with the water spout, all trials were manually blind scored with 6 categories for rewarded trials (disregard trial, no reach, groom, success, partial fail and complete fail) and 3 categories for non-rewarded trials (disregard trial, no reach and unrewarded reach). Trials which were disregarded included trials where there was either an electronic failure (e.g. water solenoid delivered too much or too little water, truncated behavioral and/or cortical activity imaging video was recorded, etc.), experimenter intervention (e.g. during training, after the animal would cause the position of the water spout to move due to vigorous grooming/reaching, etc.) or when the trial was deemed too difficult to score. No reach trials referred to those wherein the mouse did not groom or lift either both or one paw off of the resting platform in a forward reaching motion. Groom trials referred to the mouse engaging in natural grooming behavior. Complete fail trials consisted of the mouse reaching forward but being unable to make contact with or obtain the water drop. Partial fail trials consisted of the mouse reaching forward and making contact with the water drop but then being unable to bring the water to its mouth to drink. No rewarded trials scored with the category ‘unrewarded reach’ referred to either the animal engaging in grooming behavior or reaching behavior even when no water was present on the water spout. Grooming behavior was included in this category to reduce scoring subjectivity between natural grooming behavior and groom-to-reach behavior. Mice were also observed to switch between grooming and reaching the spout.

### Accelerating rotarod test

Mice were tested as previously described (Woodard et al., 2021). Briefly, mice were tested for 4 consecutive days on the rotarod (Ugo Basille) accelerated from 5 to 40 RPM over a total time period of 300 s. Mice received 3 trials per day with a 2 hr inter-trial interval (ITI). A fall was defined as the mouse falling from the rotarod or completing a rotation holding onto the rod and not trying to right itself at any point during the rotation. If a fall or full rotation occurred, the trial was ended and the time recorded. Mice that reached the maximum allowed time were scored as 300 s and the trial ended. The average latency to fall for the 3 trials was scored.

### Tapered beam test

Mice were tested using an automated touch sensing tapered beam test (Ardesch et al., 2017). Briefly, conductive paint surfaces serving as input electrodes to four 12-channel capacitive touch sensors (Adafruit Industries, New York, NY, USA) connected to a Raspberry Pi single-board computer recorded the start and finish times to traverse the beam using a custom Python script. The beam measured 100 cm in length tapering from 3.5 cm to 0.5 cm with a wider 1 cm base component extending to the left and right 1 cm below the upper surface of the beam. Mice received 4 trials per day for 5 and 7 consecutive days during the first and second round of tapered beam testing. Average time required to traverse the beam across the 4 trials was scored.

### Immunohistochemistry

Coronal brain sections were cut on a vibratome at 50 µm thickness (Leica VT1000S, Leica microsystems GmbH). Slides were then boiled in sodium citrate (10 mM sodium citrate, 0.05% Tween20, pH 6) to allow antigen retrieval. After washing, slices were permeabilized with 0.3% Triton X-100, blocked with BlockAid Blocking Solution (Molecular Probes) and Image-iT FX Signal Enhancer (Molecular Probes) before 1:100 primary antibody labeling overnight (rabbit monoclonal anti-DARPP32 (Abcam; ab40801) and mouse anti-NeuN (MilliporeSigma; ZMS377)). Phosphate-buffered saline, 0.1% Tween 20 (PBST) washing was then followed by 1:1000 secondary antibody labeling (rhodamine (TRITC) AffiniPure goat anti-mouse IgG(H+L) (JacksonImmuno Research Laboratories Inc.; 115-025-003) and AlexaFluorTM647-R-phycoerythtin goat anti-rabbit IgG(H+L) (A20991,Thermo Fisher)). Sections were washed with PBST and mounted on glass coverslips with Prolong™ Gold Antifade Mountant (Thermo Fisher; P36930) for subsequent imaging. Sections were imaged with a 10x and 63x objective using an up-right Leica imaging system (SP8 DIVE). Staining intensity was determined using ImageJ software. Relative intensity values are expressed relative to background.

### Kinematic and mesoscale GCaMP analysis

To accommodate for varying rewarded trial lengths, the first 10 s were examined for all rewarded and non-rewarded trials.

#### Kinematic analysis of forelimb skilled reaching behavior

Deeplabcut as described in (Mathis et al., 2018) was used to track body parts (right and left forepaws and mouth) and equipment landmarks of interest (platform and spout). Subsequent analyses were conducted using a custom Matlab code. The distance from the height of the platform to the height of the spout was calculated and represents the distance to the spout (spout distance). The euclidean distance of the left paw trajectories was calculated from the time of water reward delivery to 1.1 s afterwards for all successful rewarded trials. This time period was selected since it corresponded to the time needed to complete a successful reach. Euclidean distances traveled by the left paw were then binned. Histogram bin size reflects multiples of spout distance. For example, bin 4 contains successful rewarded trials where the euclidean distances of the left paw trajectories were 4x that of the spout distance. Euclidean distances are reported as multiples of spout distance since this distance represents the most efficient route the left paw could take to reach the spout. The average euclidean distance and standard deviation were calculated for each mouse then genotype averaged.

#### GCaMP image processing and analysis

All GCaMP image processing and analysis were conducted using custom Matlab codes. All GCaMP responses were movement and hemodynamic artifact corrected by subtracting changes in green reflectance signals from observed green epi-fluorescence (Vanni et al., 2017) and expressed as percentages relative to baseline responses (F-F_0_/F_0_)*100 where F_0_ is the baseline from the start of the trial to water reward delivery. For region-based analysis, the brain-to-atlas approach in MesoNet (Xiao et al., 2021) was used to register cortical images to a common atlas using predicted cortical landmarks to determine regions of interest (ROIs). A 5 x 5 pixel region centered in each ROI was used for examination of peak amplitude and baseline standard deviation. Peak amplitude was calculated from the baseline (defined as 1-5 s from the start of the trial) to the peak. Cortical area activated was determined as pixel intensities greater than 4x standard deviation of the baseline (1-5 s). Contralateral (right) and ipsilateral (left) hemisphere ROIs include the primary motor (M1), secondary motor (M2), somatosensory mouth (sspm), somatosensory forelimb (sspfl), somatosensory hindlimb (ssphl), somatosensory area unassigned (sspun), somatosensory nose (sspn), somatosensory barrel field domain (sspbfd), somatosensory trunk (ssptr), primary visual (visp), retrosplenial lateral agranular part (rspagl) and retrosplenial dorsal (rspd) cortices. Examination of the ΔF/F standard deviation during time windows before the visual cue and reward revealed no differences between genotypes (data not shown). As such, subsequent analysis was concentrated during the time period after the water reward (until 4 s afterwards).

### Experimental design, statistical analysis and code accessibility

All experimenters were blinded during the analysis. Unless otherwise stated, Two-way Anova and Šídák’s multiple comparisons post-hoc test were used. Statistical analysis was calculated using Graphpad Prism. Alpha level for all tests was p=0.05. The code used for the analysis is available from the corresponding authors upon request.

## Results

### Overview of experimental assessment timeline

The behavioral testing timeline is depicted in Figure 1A. After ∼1 week of chronic window and head-fixation bar surgical recovery, all mice underwent tapered beam training and testing to examine baseline gross motor function. Mice were then water restricted and trained to perform skilled forelimb water-reaching (Fig. 1B). Behavioral camera recording combined with markerless pose estimation (Mathis et al., 2018) enabled tracking and assessment of progressive forelimb coordination defects. Simultaneous recording of cortical activity using GCaMP6 mesoscale imaging further enabled assessment of progressive cortical circuit changes. After the completion of water-reaching assessment, mice were allowed to rest and readjust to *ad libitum* water consumption. Mice then underwent rotarod testing and a second round of tapered beam testing to confirm HD motor defects seen in the water-reaching task. Finally, immunohistochemistry staining of DARPP-32, a striatal medium spiny neuron (MSN) marker, was conducted to confirm HD pathology. Mice were weighed daily to monitor health (data not shown).

**Figure 1:**
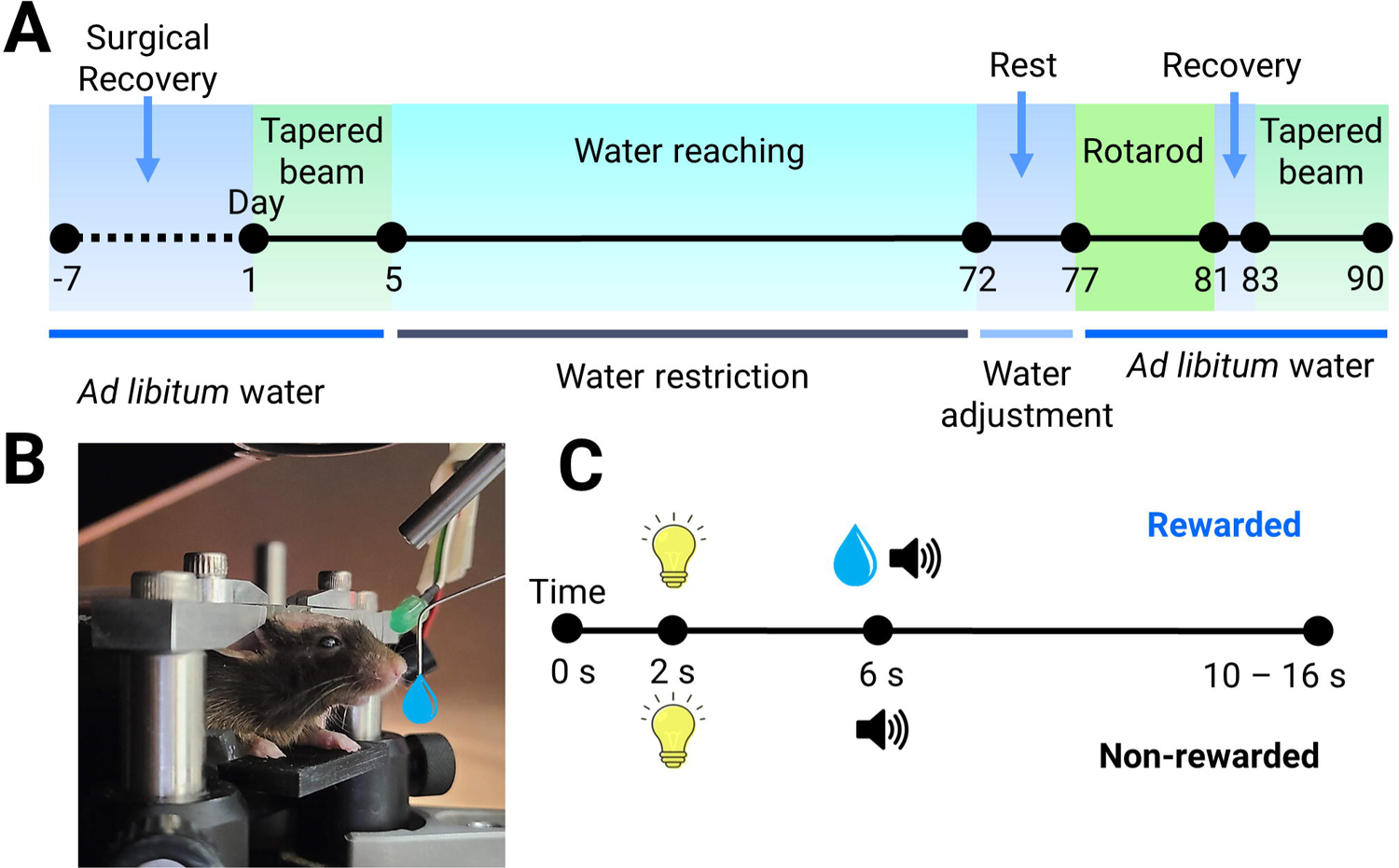
Scheme of behavioral testing. **(A)** Animals were allowed to recover from surgery (∼1 week) before initial tapered beam testing (5 days) which was followed by water-restricted forelimb water-reaching testing (∼67 days). Mice were then readjusted to *ad libitum* water consumption (∼5 days) before rotarod testing (4 days) and final tapered beam testing (7 days) with one day in between rotarod and tapered beam testing to allow stamina recovery. **(B)** Side-view image of a representative head-fixed mouse in the water-reaching task. **(C)** Trial structure for the water-reaching task where alternating rewarded and non-rewarded trials were performed. The intertrial interval was 4 s. A visual cue 2 s after initiation of camera recording was followed by an auditory cue and water drop reward for rewarded trials and only an auditory cue for non-rewarded trials both 6 s after initiation of camera recording. If a spout touch was detected after water reward delivery, the rewarded trial ended 4 s after the spout touch was detected. In cases where a spout touch was not detected, the rewarded trial timed out 10 s after water reward delivery. All non-rewarded trials ended 10 s after the auditory cue. Total trial length was therefore 16 s for non-rewarded trials and could range from 10-16 s for rewarded trials depending on if a water spout touch was detected. For consistency, all trial lengths were truncated to 10 s in total for subsequent analyses.

### Progressively reduced forelimb motor performance in HD mice

Trial structure for the water-reaching task with alternating rewarded and non-rewarded trials is depicted in Figure 1C. Briefly, a visual cue was delivered 2 s after initiation of camera recording for all trials. For rewarded trials, an auditory cue and water drop was delivered 4 s later (6 s since start of trial). For non-rewarded trials only an auditory cue was given with no water delivered. Water-reaching performance in both genotypes was quantified over 60 days (Fig. 2). Over time mice were trained during rewarded trials to reach forward towards the spout from their resting position (reach-to-grasp behavior), grasp the water drop then successfully bring the water to their mouth to drink (grasp-to-drink behavior)(Fig. 3A)(Supplemental Video 1-2). We refer to this overall as the ‘*reach-grasp-drink*’ movement. Although no aversive punishment was given for reaching the spout during non-rewarded trials, no water reward was available on the spout making any attempts futile.

**Figure 2:**
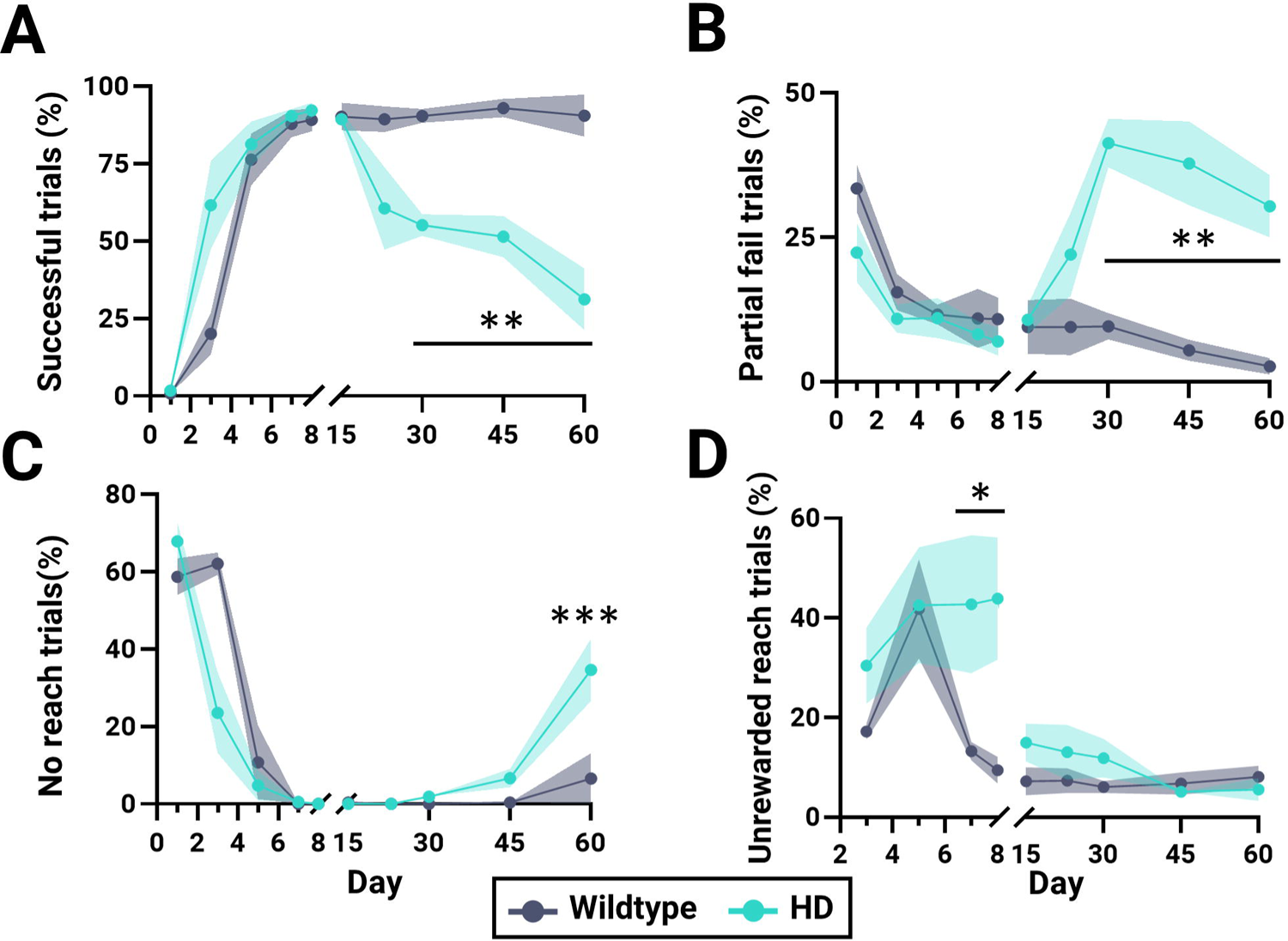
Water-reaching task behavioral categorization. HD (n = 6) and WT (n = 4) mice are denoted in teal and gray, respectively. **(A-C)** Percent of successful. **(A)**, partial fail **(B)** and no reach **(C)** trials to the total number of rewarded trials over time. **(D)** Percent of unrewarded reach trials (reaching occurs despite there being no reward) to the total number of non-rewarded trials over time. Shaded intervals denote standard error of the mean. ***, ** and * denotes p <0.005, <0.01 and <0.05, respectively.

**Figure 3:**
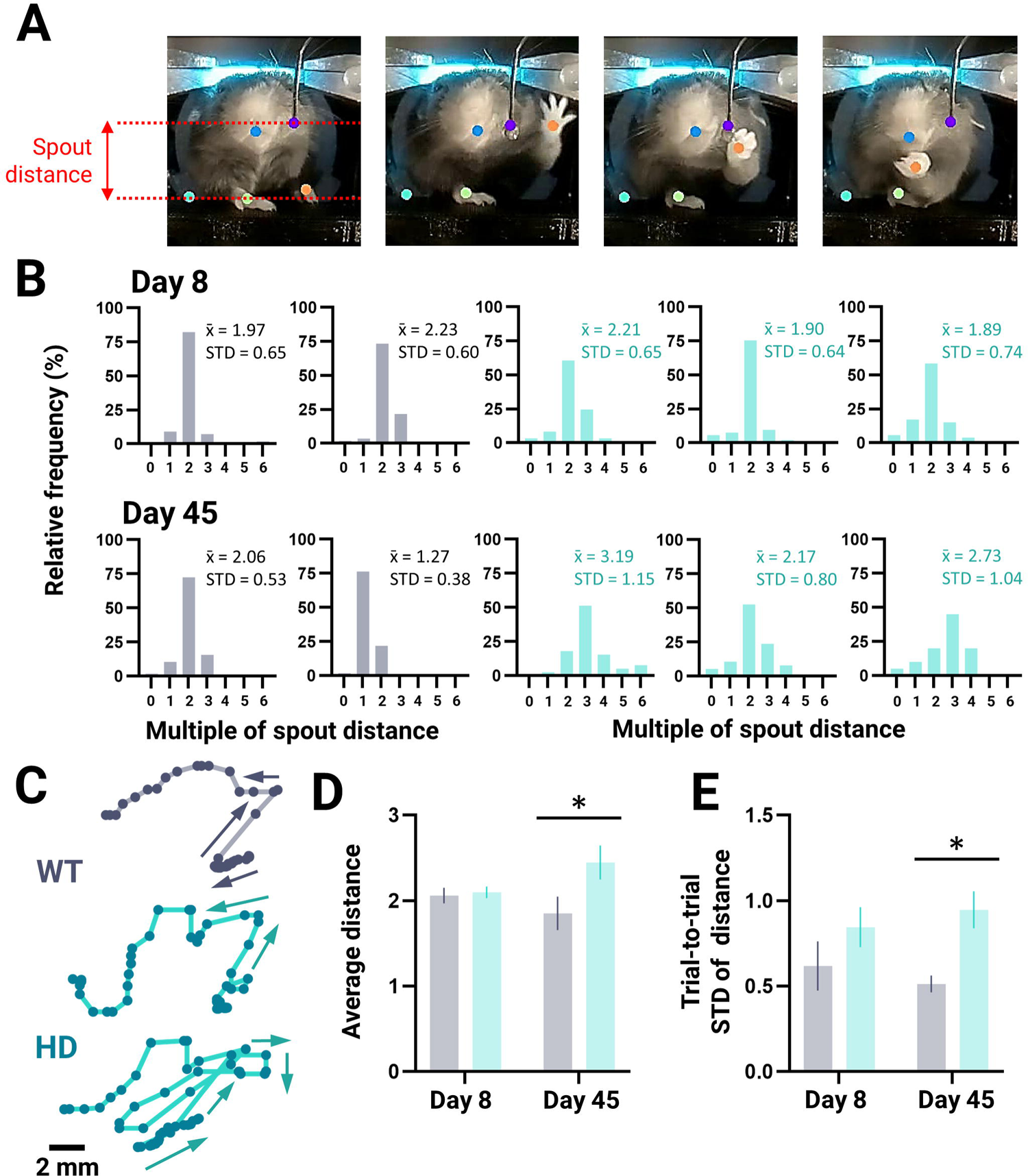
Kinematic analysis of successful trials. HD (n = 6) and WT (n = 4) mice are denoted in teal and gray, respectively. **(A)** Representative images depicting the mouse at rest and the reach-grasp-drink water-reaching movement. Dots represent either different body parts or equipment labeled for use in markerless pose estimation. The spout distance (calculated from the height of the platform to the height of the spout; see Methods for more details) is depicted in red. **(B)** Distribution of euclidean distance traveled (water reward delivery to 1.1 s afterwards) by the left paw during successful rewarded trials on Day 8 (top graphs) and Day 45 (bottom graphs) for representative WT and HD mice. The distance traveled in each trial was binned with intervals reflecting how many more times the path taken was compared to the spout distance (see Methods for more details). Relative frequencies (%) of each bin are reported. Average euclidean distance traveled (x^-^) and standard deviation (STD) are indicated and reflect multiples of spout distance. **(C)** Representative left paw X, Y trajectories of WT (top trace) and HD (middle and bottom traces) mice performing a successful water-reaching movement during a rewarded trial on Day 45. Arrows denote path of trajectory. **(D-E)** Average euclidean distance traveled across all successful trials **(D)** and trial-to-trial variability (Standard deviation; STD) of successful reaching trajectories **(E)** on Day 8 and 45. Measurements are given in multiples of spout distance; D and E are per mouse averages. * denotes p<0.05.

On Day 1 there were minimal successful trials as both groups were learning the task (Fig. 2A). In both genotypes (Day 1) the largest proportion of rewarded trials were spent not engaging with the task (no reach)(WT: 58.7 ± 4.7%; HD: 67.9 ± 5.0)(Fig. 2C) with the second largest proportion of trials spent reaching the water spout to swat the water away (partial fail)(WT: 22.3 ± 5.1%; HD: 33.4 ± 4.2%)(Fig. 2B). After performing successful trials for the first time on Day 3 (20.1 ± 6.8%)(Fig. 2A), WT mice engaged in frequent unrewarded reaching and/or groom-to-reach behavior on Day 5 (41.8 ± 10.0%)(Fig. 2D). WT mice however, quickly decreased unrewarded reaching behavior and by Day 8 performed minimal unrewarded reach trials (9.4 ± 2.7%). WT mice achieved a near perfect success performance rate by Day 8 (WT: 89.2 ± 3.7%)(Fig. 2A).

Similar to WT mice, HD mice also after performing successful trials for the first time on Day 3 (61.6 ± 14.3%)(Fig. 2A) engaged in frequent unrewarded reaching and/or groom-to-reach behavior on Day 5 (42.6 ± 11.6%)(Fig. 2D). The frequency of unrewarded reaching however, persisted. Although HD mice also reached near perfect success performance rates by Day 8 (HD 92.3 ± 2.5%)(Fig. 2A) significantly more unrewarded reaching was still present on this day compared to WT (WT: 9.4 ± 2.7%; HD: 43.9 ± 12.3%; p=0.0169). By Day 15, the unrewarded reach trials in HD mice decreased to the same frequency as seen in WT mice.

Over time WT mice were able to maintain their high success rate until at least Day 60 (Fig. 2A). HD mice however, experienced a progressive decline in successful trials. By Day 60 the successful performance rate was 31.1 ± 10.0% for HD mice. No significant changes in weight were seen throughout the entire behavioral testing timeline indicating water restriction was not the cause of HD mice performance decline (data not shown).

Unsuccessful trials were divided and scored as either no reach, groom, partial fail and complete fail trials (Extended Data Fig. 2-1). Partial fail scores denote trials where the mouse made contact with the spout and removed the water drop from the spout but was unable to retain the water drop (successful reach-to-grasp performance but failed grasp-to-drink performance). Complete fail scores denote trials where the mouse lifted their paw in a reaching behavior but the paw did not make contact with the spout (failed reach-to grasp performance).

The low prevalence of complete fail trials for the duration of behavioral testing for HD mice (Extended Data Fig. 2-1A) suggest that failure to perform the reach-to-grasp segment of the forelimb movement does not explain the decline in successful performance. HD mice, however, develop a significant increase in partial fail trials compared to WT mice by Day 30 (WT: 9.6 ± 2.2%; HD: 41.3 ± 4.2%; p=0.0024)(Fig. 2B) suggesting instead the grasp-to-drink segment of the movement was impaired. By Day 60 HD mice also developed a significant increase in no reach trials compared to WT mice (WT: 6.5 ± 6.5%; HD: 34.6 ± 8.1%; p=0.0007)(Fig. 2C). Throughout the whole duration of behavioral testing, mice in both genotypes spent a minimal number of rewarded trials grooming (Extended Data Fig. 2-1).

### Increased distance and variable forelimb reaching movement in HD mice

On average, WT mice were able to obtain the water drop 1.0 ± 0.1 s after reward delivery. As such, markerless pose estimation was used to track the left paw from the time of water reward delivery to 1.1 s afterwards (Fig. 3). Euclidean distance traveled by the left paw during successful trials is presented as multiples of the spout distance over this fixed period of time. Sample paired distribution of reaching trajectory distances for two WT (gray) and three HD (teal) mice on Day 8 (top panels) and Day 45 (bottom panels) with corresponding average euclidean distance and trial-to-trial standard deviation are shown (Fig. 3B)(all mice are shown in Extended Data Fig. 3-1). Unlike on Day 8 when the average euclidean distance traveled by the left paw was the same in both genotypes (WT: 2.1 ± 0.1; HD: 2.1 ± 0.1 spout distances; p=0.9811), the left paw of HD mice traveled a greater distance during the reach on Day 45 than WT mice (WT: 1.9 ± 0.2; HD: 2.4 ± 0.2 spout distances; p=0.0320)(Fig. 3D). Sample left paw reaching trajectories on Day 45 are shown for WT and HD mice (Fig. 3C). The variability in reaching distances for all successful trials on Day 8 (measured as the standard deviation) was not statistically different between genotypes (WT: 0.62 ± 0.14; HD: 0.84 ± 0.12 spout distances; p=0.3397)(Fig. 3E). On Day 45 however, HD mice displayed a greater variability in reaching trajectory distances than WT mice (WT: 0.51 ± 0.05; HD: 0.95 ± 0.11; p=0.0360).

### Changes in cortical activity dynamics during reaching over time in HD mice

The brain-to-atlas approach in MesoNet (Xiao et al., 2021) was used to register cortical images to a common atlas using predicted cortical landmarks. Regions of interest (ROIs) were then defined (Fig. 4A; ROIs are color and number labeled). Sample heat maps of trial-to-trial cortical ΔF/F for select ROIs during successful, unrewarded reach and/or partial fail trials on Day 8 and/or 45 from a WT and HD mouse are shown (Fig. 4B)(all mice are shown in Extended Data Fig. 4-1 to 4-3).

**Figure 4:**
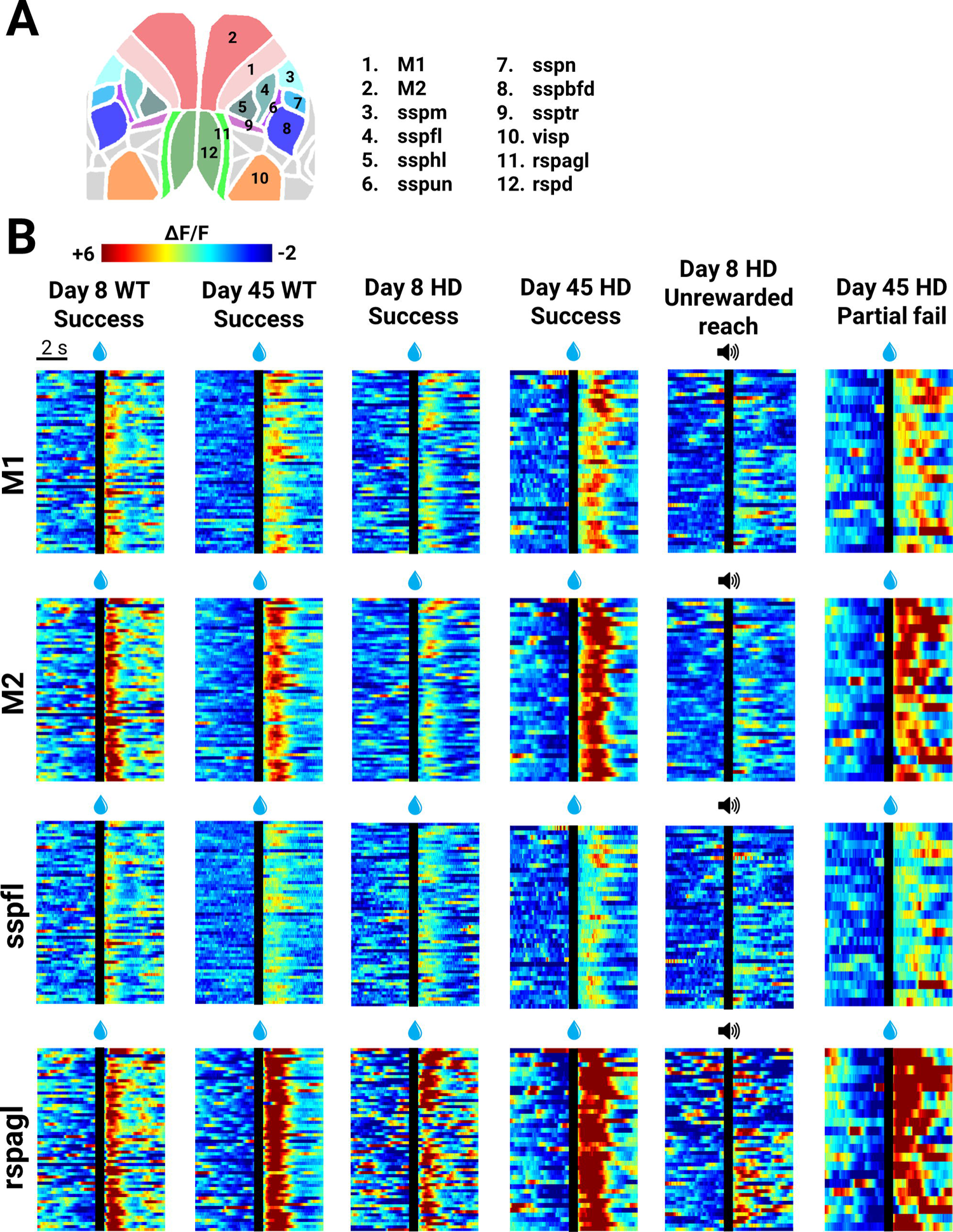
Representative trial-to-trial GCaMP cortical activity in regions of interest. **(A)** Cartoon depicts regions of interest investigated in subsequent analyses. **(B)** Representative trial-to-trial heat-map of GCaMP (ΔF/F) cortical activity in contralateral M1 (primary motor), M2 (secondary motor), sspfl (somatosensory forelimb) and rspagl (retrosplenial lateral agranular) from a WT and HD mouse on Day 8 and Day 45 for success, unrewarded reach and/or partial fail trials. Individual trials are stacked in rows. Time of the water reward (for rewarded trials) and tone (for non-rewarded trials) is denoted with a black line.

Sample time series of GCaMP6 cortical wide-field imaging on Day 8 and 45 are shown in the top panels of Figure 5A for a representative WT and HD mouse. On Day 8 despite both HD and WT mice having comparable success rates (Fig. 2A) and reaching distances (Fig. 3D), genotype differences in cortical activity were apparent (Extended Data Fig. 5-1). Examining specific ROIs further revealed that the peak amplitude across all ROIs in both the contralateral and ipsilateral hemisphere was greater in WT compared to HD mice (contralateral: F_1,8_=5.925, p=0.0409, ANOVA; ipsilateral: F_1,8_=5.967, p=0.0404, ANOVA)(Extended Data Fig. 5-1). Together this indicates that on Day 8, more extensive cortical activation associated with reaching was seen in WT compared to HD mice.

**Figure 5:**
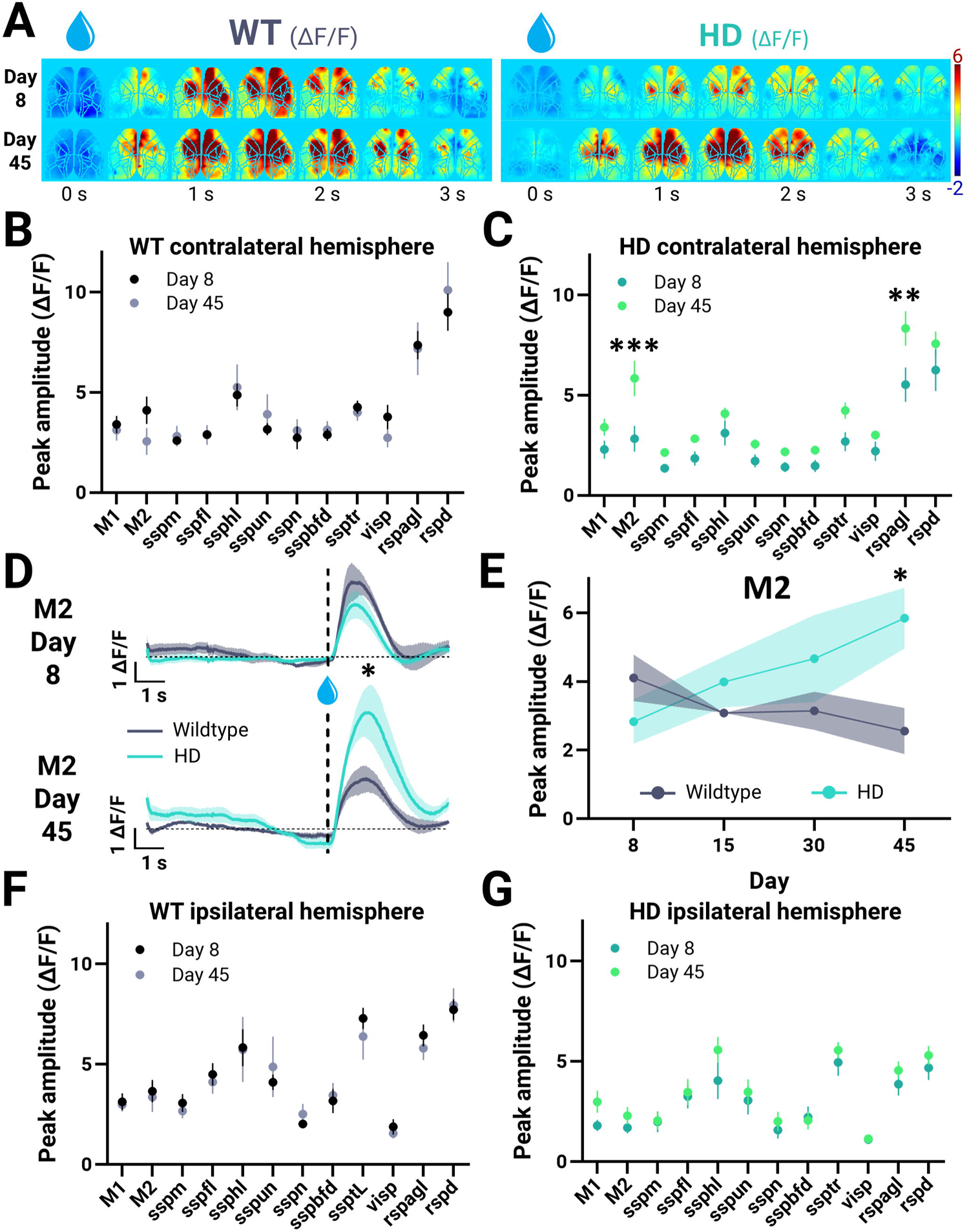
Longitudinal mesoscale GCaMP imaging of the cortex during water-reaching. HD (n = 6) and WT (n = 4) mice are denoted in teal and gray, respectively. 5×5 pixel regions are centered in regions of interest (ROIs) and examined. **(A)** Time series of cortical wide-field GCaMP imaging (ΔF/F) from a representative WT (left panels) and HD (right panels) mouse on Day 8 (top panels) and Day 45 (bottom panels) during successful trials. **(B-C)** Peak ΔF/F amplitude of ROIs in the contralateral hemisphere on Day 8 and Day 45 for WT (F_1,72_=0.006, p=0.9388, ANOVA)**(B)** and HD (F_1,10_=5.521, p=0.0407, ANOVA)**(C)** mice. **(D)** Time course of M2 (secondary motor cortex) activation on Day 8 (top panel) and Day 45 (bottom panel). Vertical dotted line denotes time of water reward delivery. Horizontal dotted line denotes zero ΔF/F level. Significance reflects a difference in genotype peak response. **(E)** Corresponding change in peak ΔF/F amplitude of M2 over time. **(F-G)** Peak ΔF/F amplitude of ROIs in the ipsilateral hemisphere on Day 8 and Day 45 for WT (F_1,6_=0.026, p=0.8770, ANOVA)**(F)** and HD (F_1,10_=0.709, p=0.4193, ANOVA)**(G)** mice (WT and HD day factor: not statistically significant) ***, ** and * denotes p<0.005, <0.01 and <0.05, respectively.

When examined longitudinally within genotypes (comparison of Day 8 to Day 45), the peak amplitude across all ROIs in the contralateral hemisphere increased in HD mice (F_1,10_=5.521, p=0.0407, ANOVA)(Fig. 5C) with no significant changes in WT mice (F_1,72_=0.006, p=0.9388, ANOVA)(Fig. 5B). In particular, the pixel regions centered in the contralateral secondary motor cortex (M2) and retrosplenial cortex lateral agranular part (rspagl) displayed significantly greater peak amplitude over time in HD mice (Fig. 5C). No significant changes in peak amplitude across all ROIs in the ipsilateral hemisphere were seen over time for WT (F_1,6_=0.026, p=0.8770, ANOVA) and HD (F_1,10_=0.709, p=0.4193, ANOVA) mice when comparing within genotypes (Fig. 5F-G).

When comparing genotypes (WT to HD) a significant difference in contralateral M2 peak activity was seen on Day 45 (WT: 2.6 ± 0.7; HD: 5.8 ± 0.9; p=0.0455)(Fig. 5E). Figure 5D shows the average time course of contralateral M2 ΔF/F activation on Day 8 and 45 for all mice. Genotype differences were also seen in contralateral sspm, sspbfd and visp cortices on Day 8 (Extended Data Fig. 5-2). Comparing WT to HD mice overtime further reveals differences in ipsilateral M2, rspagl and retrosplenial cortex, dorsal part (rspd)(Extended Data Fig. 5-3).

### Unrewarded reach, fail vs success trials performed by HD mice

In addition to performing progressively increased fail trials over time (Fig. 2B), HD mice also engaged in unrewarded reaching behavior for more days (Fig. 2D). We examined the cortical activity underlying these trial types further and compared them to successful trials. Sample time series of cortical imaging on Day 8 from a representative HD mouse is shown in Figure 6A for all successful, failed and unrewarded reach trials. On Day 8, compared to successful and failed trials, the peak amplitude of all ROIs in the contralateral and ipsilateral hemisphere was significantly reduced in unrewarded reach trials (Fig. 6B-C). In particular, the peak cortical activity at the contralateral rspd and rspagl was significantly reduced in unrewarded trials compared to successful and failed trials. Figure 6D shows the average time course of contralateral rspagl and rspd ΔF/F activation for all three trial types and mice on Day 8. A significantly greater area was also activated after the water reward when HD mice were performing successful and failed trials than during unrewarded reach trials (Fig. 6E). No differences in peak amplitude or area activated were seen when comparing successful to failed trials on Day 45 (Extended Data Fig. 6-1).

**Figure 6:**
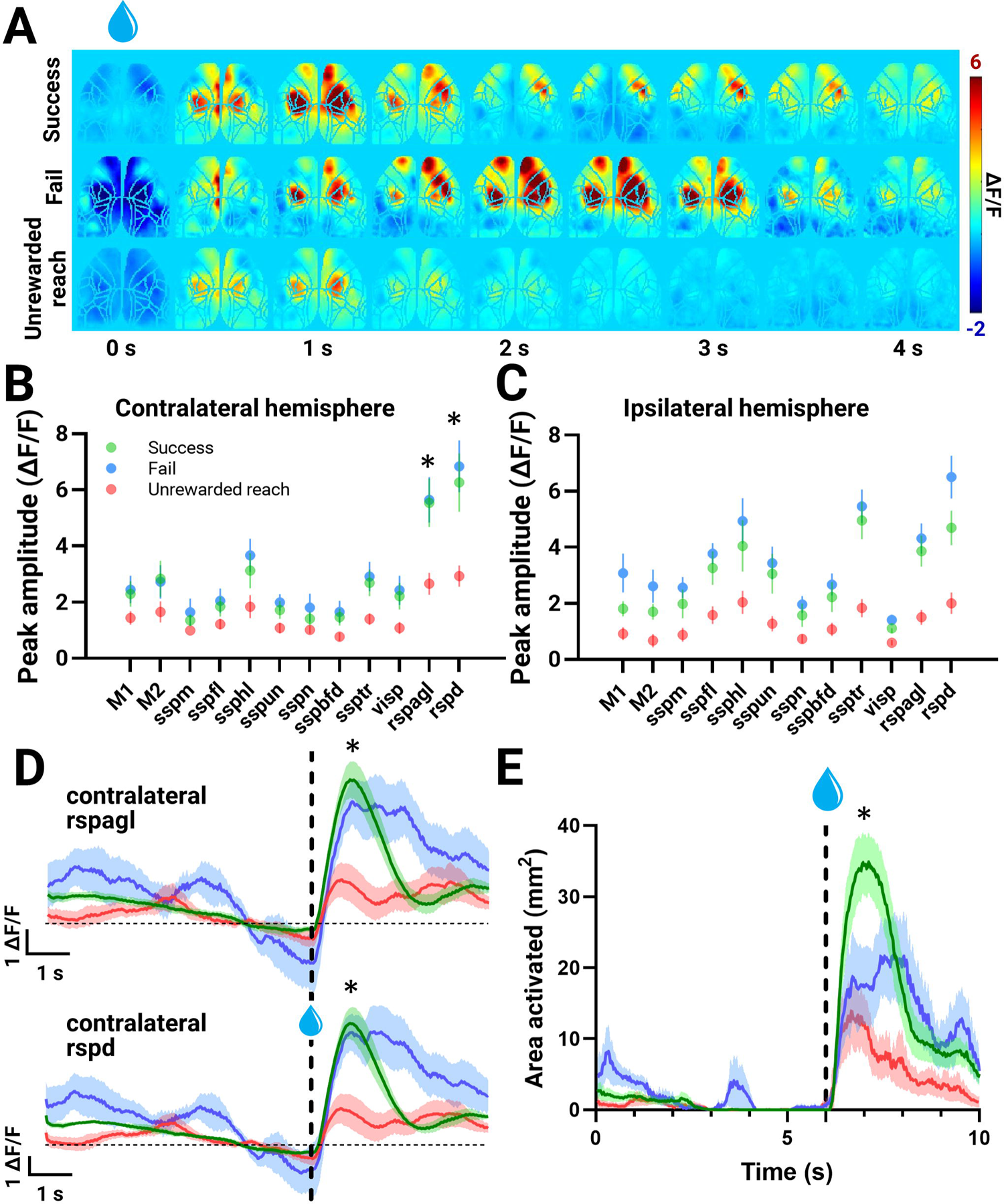
Mesoscale GCaMP imaging of the cortex during success, fail and unrewarded reach trials performed by HD mice on Day 8. HD (n = 6) success, fail and unrewarded reach trials are denoted in green, blue and red, respectively. **(A)** Time series of cortical wide-field GCaMP imaging (ΔF/F) from a representative HD mouse on Day 8 during success (top), fail (middle) and unrewarded reach (bottom) trials. **(B-C)** Peak amplitude of regions of interest in the contralateral (F_2,15_=3.862, p=0.0444, ANOVA)**(B)** and ipsilateral (F_2,15_=9.526, p=0.0021, ANOVA)**(C)** hemisphere for different trial types. **(D)** Time course of contralateral rspagl (retrosplenial cortex lateral agranular part) and rspd (retrosplenial cortex dorsal part) activity. Vertical dotted line denotes time of water reward delivery. Horizontal dotted line denotes zero ΔF/F level. Significance reflects a difference in unrewarded reach peak response compared to other trial types. **(E)** Area activated across the entire trial duration for success, fail and unrewarded reach trials. The threshold was set at 4x standard deviation (STD) of the baseline. Significance of area activated after the water reward for unrewarded reach trials compared to other trial types is indicated. * denotes p<0.05.

### Forelimb coordination and movement defects are coincident with gross motor defects and HD pathology

To confirm the progressive aberrant forelimb movement phenotype characterized with water-reaching testing, HD mice were examined and compared to WT mice using classical tapered beam and rotarod testing (Fig. 7A-D). Before the water-reaching assessment, with the exception of Day 1 when WT mice learned to traverse the tapered beam faster than HD (WT: 5.418 ± 0.618 s; HD: 8.320 ± 0.997 s; p=0.0265), both genotypes spent on average, the same time traversing the tapered beam (Fig. 7A) indicating HD mice likely do not have a motor deficit at this stage. All mice were also subjected to a second round of tapered beam testing after water-reaching assessment (Fig. 7B). During this second testing phase, with the exception of Day 1, WT mice traversed the beam significantly faster than HD mice. We then compared the performance of the mice during the first round of testing (mice aged ∼5 months) with the second round (mice aged ∼8 months). Day 4 has previously been used as the first testing day after successful learning of the task (Ardesch et al., 2017). Comparison of Day 4 during the first round of testing to the last assessment day (Day 7) during the second round of testing, revealed that WT mice traversed the beam in a faster time by the last assessment day (first testing round Day 4: 4.728 ± 0.532 s; second testing round Day 7: 2.97 ± 0.077 s; p=0.0044)(Fig. 7C). For HD mice, the time to traverse the beam did not change between the first (Day 4: 5.541 ± 0.614 s) and second (Day 7: 5.550 ± 1.192 s) round of testing (Fig. 7C).

**Figure 7:**
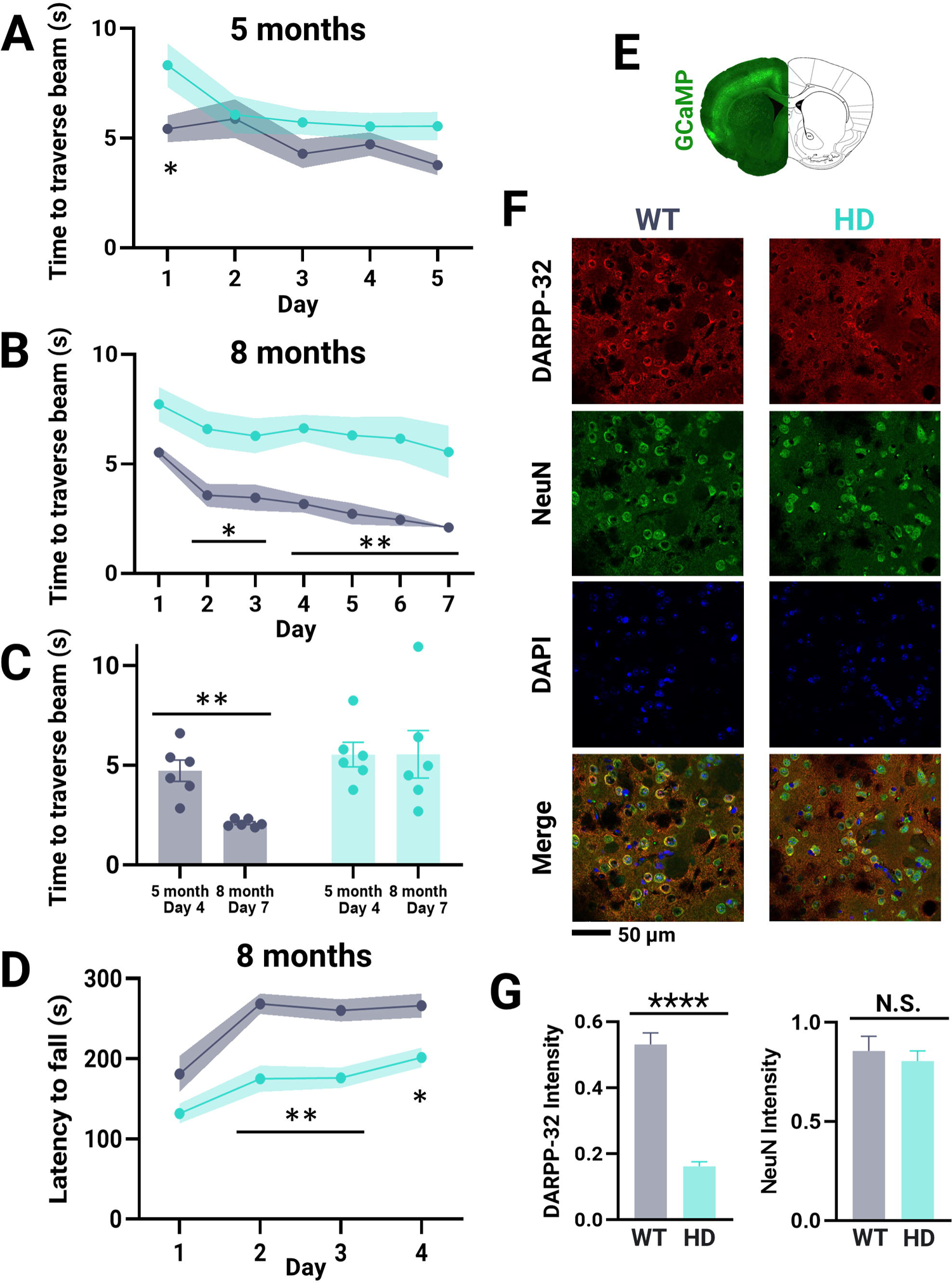
Tapered beam and rotarod gross motor assessment and post-mortem immunohistochemistry staining. HD and WT mice are denoted as teal and gray, respectively. **(A-B)** Time to traverse the tapered beam determined before (∼5 months)**(A)** and after (∼8 months)**(B)** water-reaching testing for HD (n = 6) and WT (n = 6) mice. **(C)** Time to traverse the tapered beam on the first day after completion of tapered beam learning (mice age: ∼5 month; day of testing: 4) compared to the last testing date (mice age: ∼8 month; day of testing: 7) for HD and WT mice. **(D)** Latency to fall from the rotarod determined after water-reaching testing (∼8 months) for HD (n = 6) and WT (n = 6) mice. **(E)** Representative Thy1-GCaMP6s coronal slice. DARPP-32 intensity was quantified in the striatum. **(F)** Representative images of DARPP-32, NeuN and DAPI staining in the striatum with a merged overlay from a WT and HD mouse. **(G)** DARPP-32 and NeuN intensity in the striatum of HD (n = 4) compared to WT (n = 4) mice. Error bars and shaded intervals denote standard error of the mean. ****, **, * and N.S. denote p <0.0001, <0.01, <0.05 and statistically non-significant, respectively as determined by Two-way Anova and Šídák’s multiple comparisons post-hoc test for tapered beam and rotarod tests and unpaired T-test for immunohistochemistry staining.

Further in support of the idea HD mice were experiencing motor and balance defects by the second round of tapered beam testing (∼8 months of age), HD mice also displayed impaired performance on the accelerating rotarod task as determined by a decreased latency to fall compared to WT (Fig. 7D). Together, the second round of tapered beam and rotarod testing - both performed after water-reaching assessment - suggest HD mice have reached the motor manifest stage of disease. This is consistent with the decreased success rate seen by the end of the water-reaching task in HD mice (Fig. 2A).

Finally, to confirm HD pathology, striatal MSNs which make up 95% of all neurons in the striatum were immunostained for DARPP-32 (Fig. 7E-G). Consistent with previous literature (Peng et al., 2016; Southwell et al., 2016), a significant decrease in DARPP-32 (relative intensity WT: 0.531 ± 0.035; HD: 0.162 ± 0.014; t=9.765; df=6; p<0.0001) but not NeuN (relative intensity WT: 0.86 ± 0.07; HD: 0.81 ± 0.05; t=0.5644; df=6; p=0.5930) intensity was seen in HD compared to WT mice further confirming the manifestation of HD phenotype at ∼8 months of age (Fig. 7G).

## Discussion

The shared evolutionary origin and characteristics of skilled forelimb movements (Whishaw et al., 1992; Galiñanes et al., 2018) enable translational parallels to be drawn from preclinical mouse studies. In HD patients and pre-symptomatic carriers, deficits in motor learning, temporal sequencing and coordination of voluntary movements have been reported (Feigin et al., 2006; Klein et al., 2011; Shabbott et al., 2013). Using a water-reaching task, we reveal the presence of event sequence learning defects and progressive increases in cortical activity underlying forelimb deficits in the zQ175 HD mouse model (see Fig. 8 for a summary of the results).

**Figure 8:**
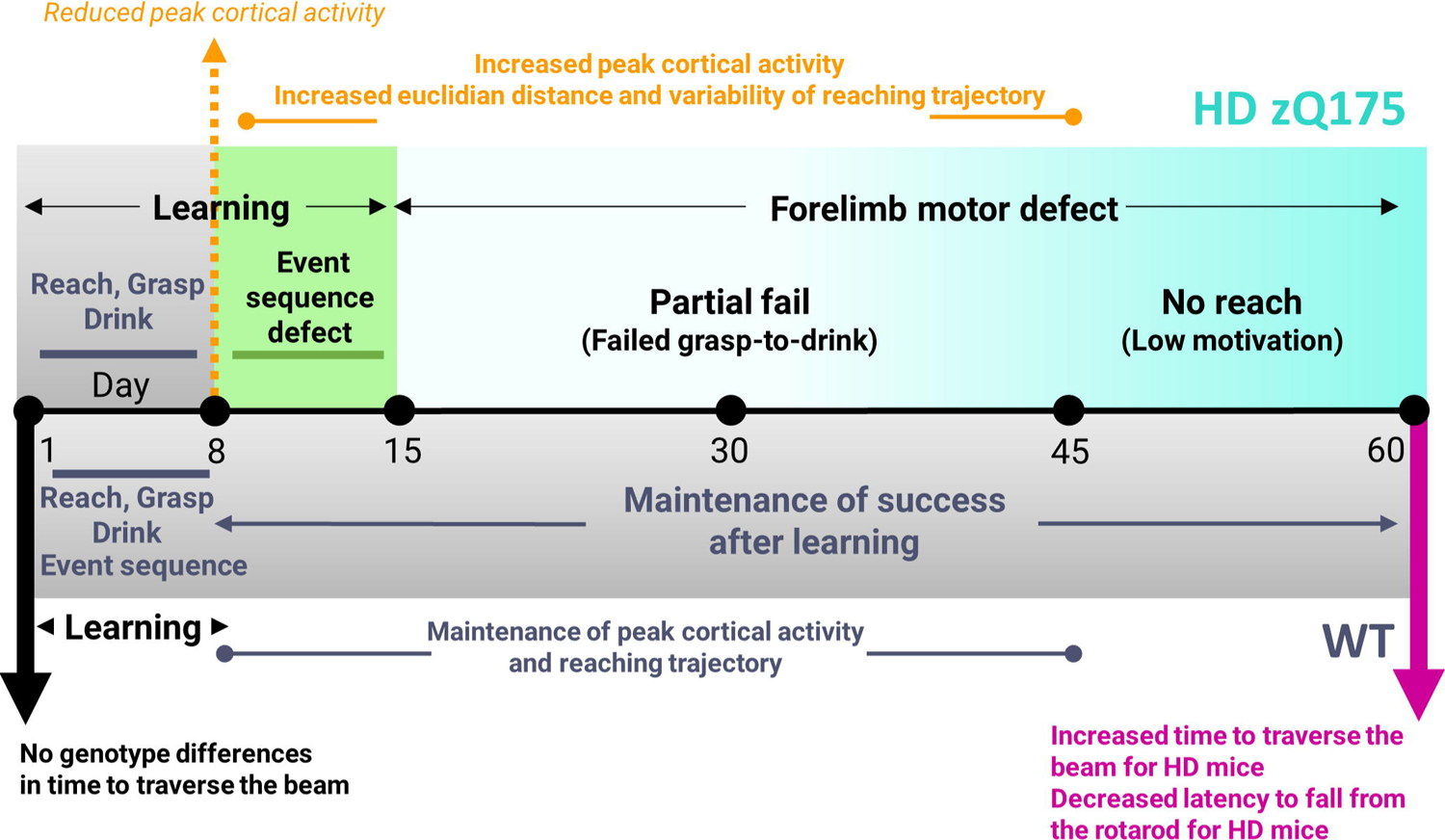
Schematic summary of altered cortical activity and motor defects in HD mice. Timeline of HD (top) and WT (bottom) learning and performance in the water-reaching task. Corresponding tapered beam and rotarod tasks used to validate the phenotypes observed in the water-reaching tasks are included. WT mice learn both the reach-grasp-drink movement and task event sequence (alternating reward then non-rewarded trial) by Day 8 (gray). Although HD mice also learn the reach-grasp-drink movement by Day 8 (gray) HD mice show reduced cortical activation compared to WT mice. HD mice also take longer to learn the task event sequence (green) than WT mice. Over time the peak cortical activity, euclidean distance and variability of the reaching trajectory increases in HD mice but little to no change was seen in WT mice. Unlike WT mice, HD mice also do not maintain their rate of successful performance overtime. HD mice experience first a progressive increase in partial fail trials then an increase in no reach trials reflecting failed grasp-to-drink then low task engagement, respectively. Overall, this indicates a progressively worsening forelimb motor coordination defect (light to dark teal) in HD mice.

### Task acquisition and performance across genotypes

For most motor tasks, initial learning is accompanied by trial-to-trial variability, enabling spatial exploration and progress towards efficient task execution (Dhawale et al., 2019). Variability is subsequently reduced after strategy formation (Churchland et al., 2006). We observe similar features since by Day 8, both HD and WT mice were able to successfully learn the reach-to-grasp movement. Although the movement was successfully learned by both genotypes, cortical activity underlying successful reaches was reduced in HD mice compared to WT. HD mice also required more days to learn the alternating reward/non-reward event sequence. We speculate that the extended continuation of reaching behavior during non-rewarded trials in HD mice could be a result of underlying cognitive defects that slow learning due to an inability to remember when to reach or failure to suppress motor movement.

Over time HD mice experienced a significant drop in successful reaches compared to WT. The increased trial-to-trial variability seen in HD mice compared to WT mice on Day 45 suggests that HD mice are attempting compensatory changes in reaching strategy at a time when they experienced a drop in performance. Consistent with this, positional error correction of the forelimb has previously been observed in consecutive reach trials (Becker et al., 2020). The significant increase in partial fail trials but not complete fail trials further suggests HD mice fail to engage in proper end-point fine motor corrections during the grasp-to-drink segment of the task (Elliott et al., 2001). Semi-flexed or closed paws have been shown to result in failed target reaching trials (Whishaw et al., 2018b) and could explain the decline in successful performance rates seen in HD mice. By Day 60, decreased task engagement was seen indicating that movement defects in HD mice increased in severity and alternative reaching strategies were no longer sufficient to mediate continued motivation and task engagement.

### Bilateral engagement of mesoscale cortical circuits during reaching

Consistent with other studies that report global activation of the cortex and involvement of the ipsilateral hemisphere during limb movement (Heming et al., 2019; Soma et al., 2019; Brunner et al., 2020; Quarta et al., 2022), our results also revealed widespread cortical activation across both hemispheres during water-reaching. Although we did not see wide-spread enhanced cortical activity in HD mice as some work indicates (Arnoux et al., 2018; Burgold et al., 2019; Sepers et al., 2021) compared to WT (except in M2), global cortical activation associated with reaching increased over time in HD mice (but not WT). The lack of increased cortical activity may be due to differences in task-performing awake versus anesthetized animals, HD mouse models and/or cortical areas examined. We speculate that this increase in cortical activity seen over time in HD mice may be driven by increases in average euclidean distance of the reaching trajectory. The increased euclidean distance seen in HD mice was a result of multiple sub-reach attempts that eventually led to successful task execution. Another explanation could involve changes in local inhibitory inputs (Cummings et al., 2009), spontaneous firing rates and/or activity of the striatum (Donzis et al., 2020) overtime during water-reaching assessment.

Examining cortical regions of interest revealed genotypic differences in peak GCaMP6 cortical responses in M2, sspm, sspbfd and visp of the contralateral hemisphere and rspagl, rspd and M2 of the ipsilateral hemisphere. Differences have been reported in tongue protrusions during freely moving pellet reaching between individual mice and trial types (Whishaw et al., 2018a). Although tongue protrusions were not evident in either WT or HD mice, adjustments to the tongue within the mouth, the mouth itself or whisking could explain the differences seen in sspm and sspbfd. Chemosensory, but not spatial or visual cues have been shown to guide water-reaching behavior (Galiñanes et al., 2018). The genotypic significance of visual areas seen in this study necessitates further investigation into visual and other cortical areas not typically examined in the context of forelimb reaching and other motor tasks.

The retrosplenial cortex also showed genotype-specific changes and has been linked to spatial memory and navigation (Czajkowski et al., 2014; Milczarek et al., 2018). Retrosplenial cortices may be involved in learning and maintaining correct spatial orientation of the paw towards the target (water reward). Further investigations are needed to fully understand the role retrosplenial cortices play in forelimb reaching and its contributions to HD phenotype.

Optogenetic cortical silencing has revealed the motor cortex is critical for the adjustment of complex grasping movements (Mohammed et al., 2020). Specifically, M2 has also been reported to encode movement distance and smoothness (Quarta et al., 2022). Our findings that HD mice fail to perform the grasp-to-drink portion of the movement (increased partial fail trials compared to WT) and have an increased average euclidean distance in their reaching trajectory compared to WT likely explains the genotypic hyperactivity seen in HD contralateral M2 compared to WT and is consistent with the previously reported roles M2 plays in forelimb reaching. This M2 hyperactivity evident at the motor manifest, but not premanifest stage is analogous to increases in striatal activity seen over time in HD (YAC128) mice during rotarod performance (Koch et al., 2022).

We acknowledge that epifluorescence wide-field calcium imaging which we use to assess excitability has reduced temporal resolution compared with voltage sensitive dyes and is sensitive to artifacts associated with light scattering, hemodynamics and movement. ROIs generated using deep learning cortical image and landmark registration to a common atlas may also not represent the same regions as those determined functionally. To mitigate some of these limitations, strobing of green reflectance light was used to correct hemodynamic artifacts (Wekselblatt et al., 2016; Vanni et al., 2017; Xiao et al., 2017). The head-fixed set-up further reduced movement. Despite some limitations, our study has demonstrated the water-reaching task can reliably characterize forelimb motor defects in HD mice and reveal aberrant cortical activity in HD mice.

At a time when HD and WT mice were capable of achieving similar success performance rates at the water-reaching task, our study and others (Peng et al., 2016; Liu et al., 2021) report no significant differences in tapered beam traverse time suggesting zQ175 mice are in the premanifest stages of HD at ∼5 months of age. Later, when HD mice experienced reduced performance at the water-reaching task, we and others also reported seeing increased time to traverse the tapered beam, decreased latency to fall from the rotarod, and decreased DARPP32 expression in striatal MSNs (Smith et al., 2014; Peng et al., 2016; Southwell et al., 2016; Liu et al., 2021) suggesting manifestation of HD motor phenotype and pathology in zQ175 mice occurs by ∼8 months of age.

Future studies could examine the contribution of diverse cortical areas (such as retrosplenial, visual and somatosensory) and subcortical regions (such as the striatum (Brunner et al., 2020), cerebellum (Guo et al., 2021) and thalamus (Sauerbrei et al., 2020)) to forelimb tasks in mouse models of HD and other movement disorders. Therapeutic rescue of the HD phenotype using optogenetics and parsing the contribution of direct-indirect striatal (Reiner et al., 1988; Albin et al., 1992; Barry et al., 2018) and M2 cortico-striatal pathways (Fernández-García et al., 2020) would also enable mechanistic understandings of HD forelimb defects. The ability of the water-reaching task to characterize HD phenotype suggests it can potentially be used to inform the onset of other movement disorders, therapeutic intervention windows and test drug efficacy.

### Multimedia

<u>Supplemental Video 1: Representative WT mouse performing a successful trial.</u> Simultaneous cortical wide-field GCaMP imaging (ΔF/F) and water-reaching behavior video (front and side view) from a WT mouse performing a successful trial. Scale bar denotes 0.5 mm.

<u>Supplemental Video 2: Representative HD mouse performing a successful trial.</u> Simultaneous cortical wide-field GCaMP imaging (ΔF/F) and water-reaching behavior video (front and side view) from a HD mouse performing a successful trial. Scale bar denotes 0.5 mm.

## Extended data figure legends

**Extended data Figure 2-1:**
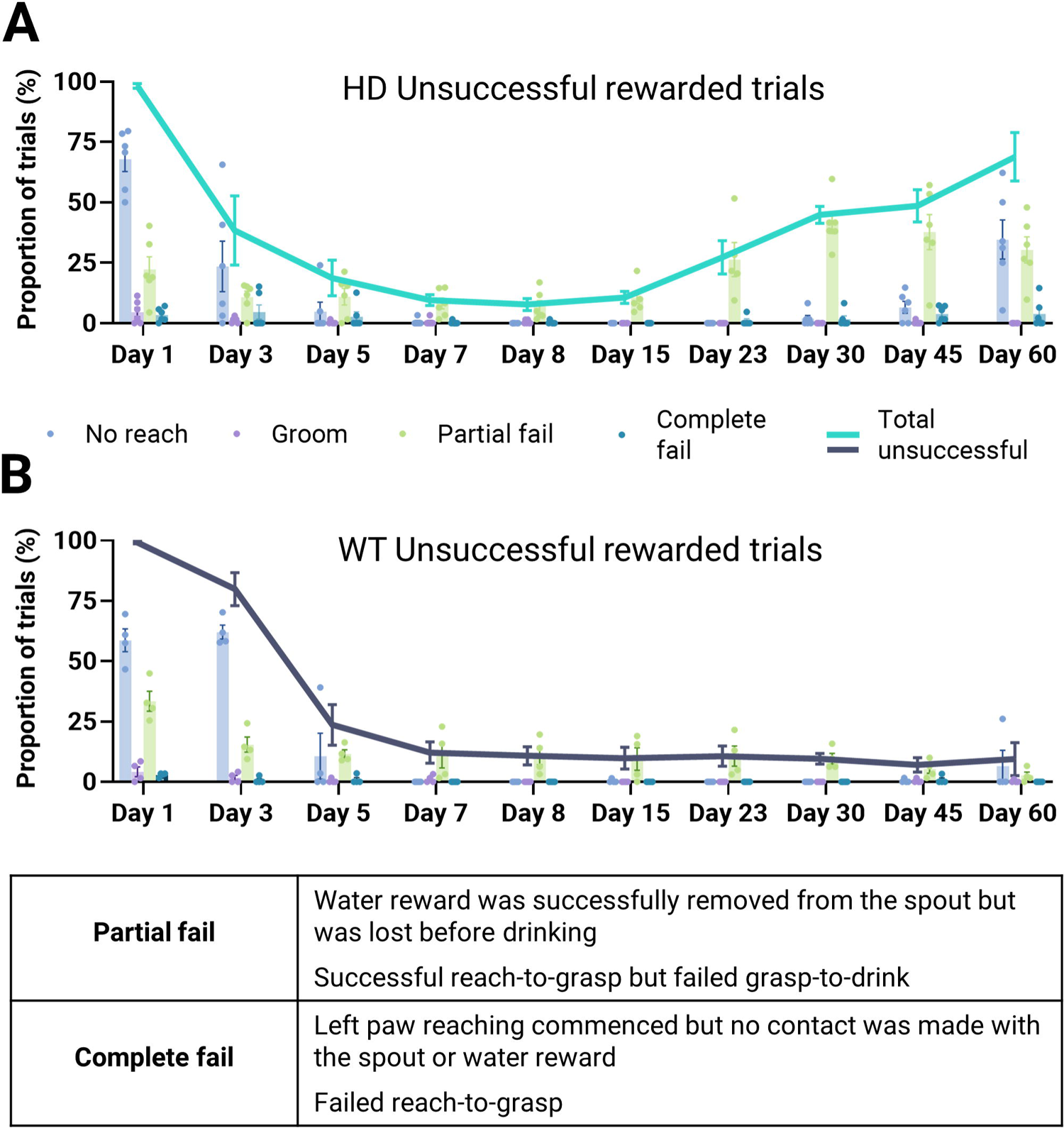
Categorization of unsuccessful rewarded trials. Proportion of unsuccessful rewarded trial types (no reach: blue; groom: purple; partial fail: green; complete fail: dark teal) and total unsuccessful trials (line) to the total number of rewarded trials for HD (n = 6)**(A)** and WT (n = 4)**(B)** mice overtime. Error bars denote standard error of the mean. Grooming and complete fail trials in both genotypes were minimal with no statistical differences between genotypes.

**Extended data Figure 3-1:**
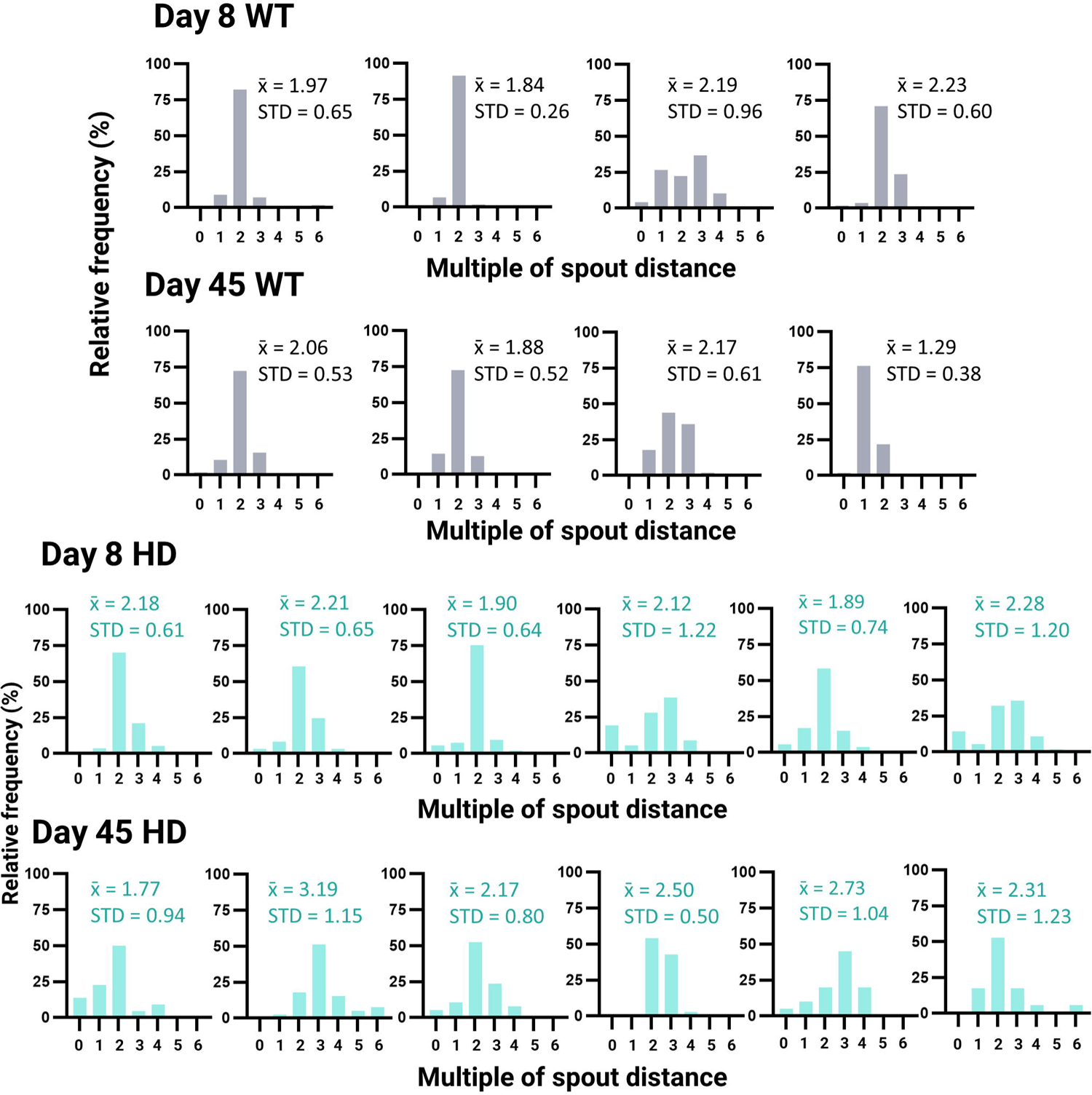
Euclidean distance distribution for successful trials on Day 8 and Day 45. Distribution of euclidean distance traveled by the left paw (water reward delivery to 1.1 s afterwards) during successful rewarded trials on Day 8 (top graphs) and Day 45 (bottom graphs) for all WT (gray) and HD (teal) mice. The distance traveled in each trial was binned with intervals reflecting how many more times the path taken was compared to the spout distance (calculated from the height of the platform to the height of the spout; see Methods for more details). Relative frequencies (%) of each bin are reported. Average euclidean distance traveled (x^-^) and standard deviation (STD) are indicated in multiples of spout distance.

**Extended data Figure 4-1:**
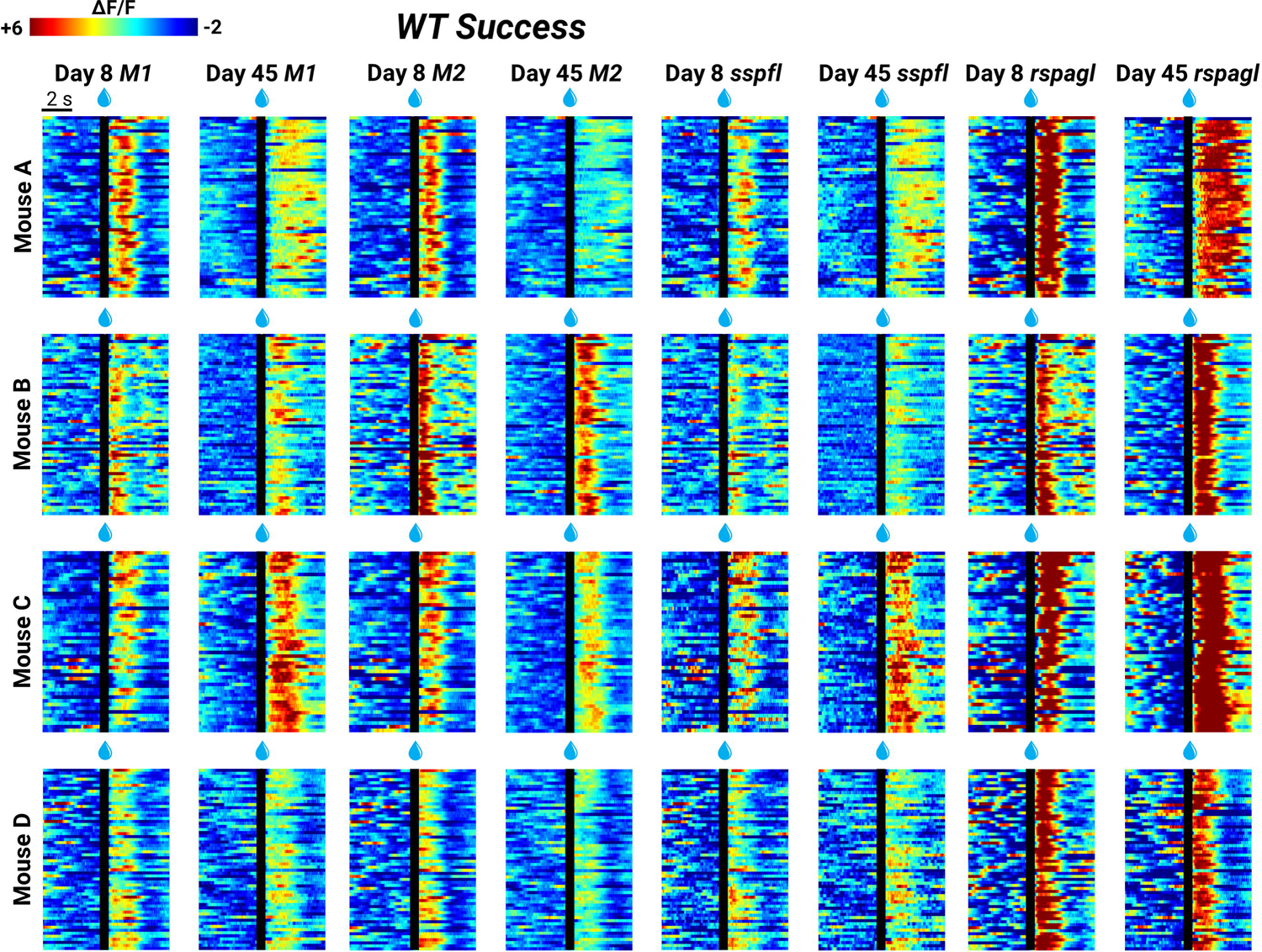
WT trial-to-trial GCaMP heat-map for successful trials. Success trial-to-trial heat-map of GCaMP (ΔF/F) cortical activity in contralateral M1 (primary motor), M2 (secondary motor), sspfl (somatosensory forelimb) and rspagl (retrosplenial lateral agranular) for all WT mice on Day 8 and Day 45. Individual trials are stacked in rows. Time of the water reward is denoted with a black line.

**Extended data Figure 4-2:**
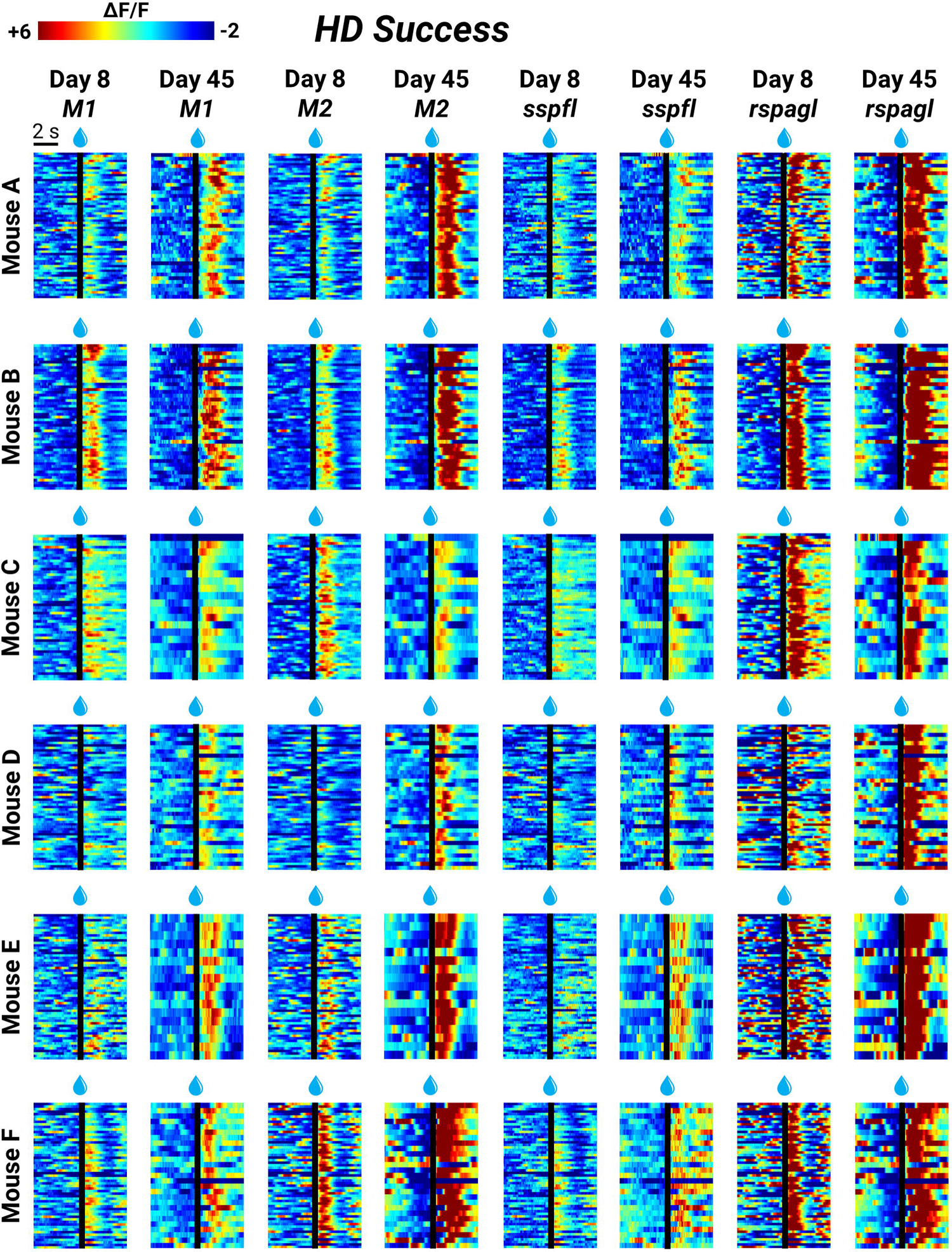
HD trial-to-trial GCaMP heat-map for successful trials. Success trial-to-trial heat-map of GCaMP (ΔF/F) cortical activity in contralateral M1 (primary motor), M2 (secondary motor), sspfl (somatosensory forelimb) and rspagl (retrosplenial lateral agranular) for all HD mice on Day 8 and Day 45. Individual trials are stacked in rows. Time of the water reward is denoted with a black line.

**Extended data Figure 4-3:**
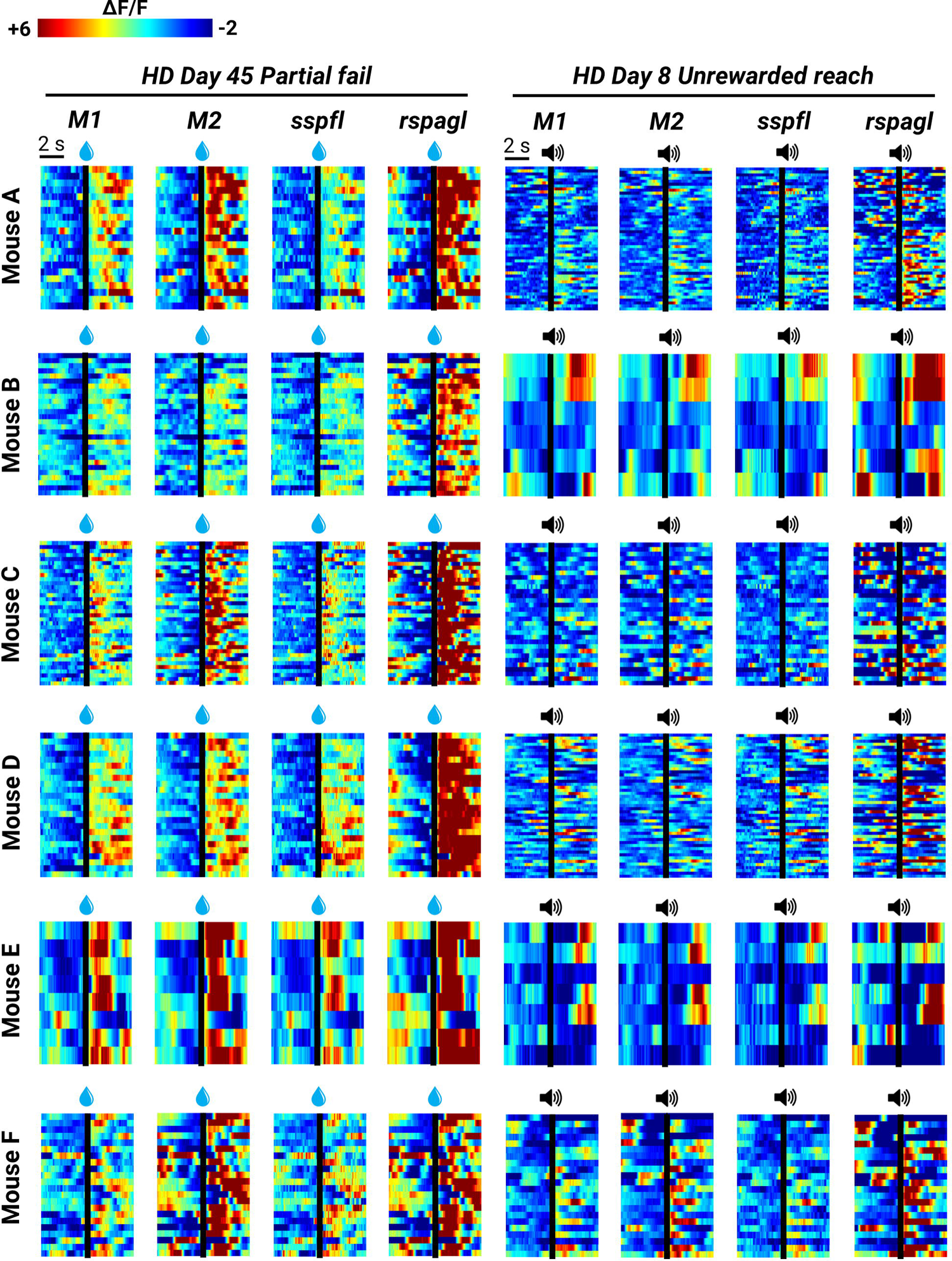
HD trial-to-trial GCaMP heat-map for unrewarded reach and partial fail trials. Partial fail and unrewarded reach trial-to-trial heat-map of GCaMP (ΔF/F) cortical activity in contralateral M1 (primary motor), M2 (secondary motor), sspfl (somatosensory forelimb) and rspagl (retrosplenial lateral agranular) for all HD mice on Day 45 and Day 8, respectively. Individual trials are stacked in rows. Time of the water reward (for partial fail trials) and tone (for unrewarded reach trials) is denoted with a black line.

**Extended data Figure 5-1:**
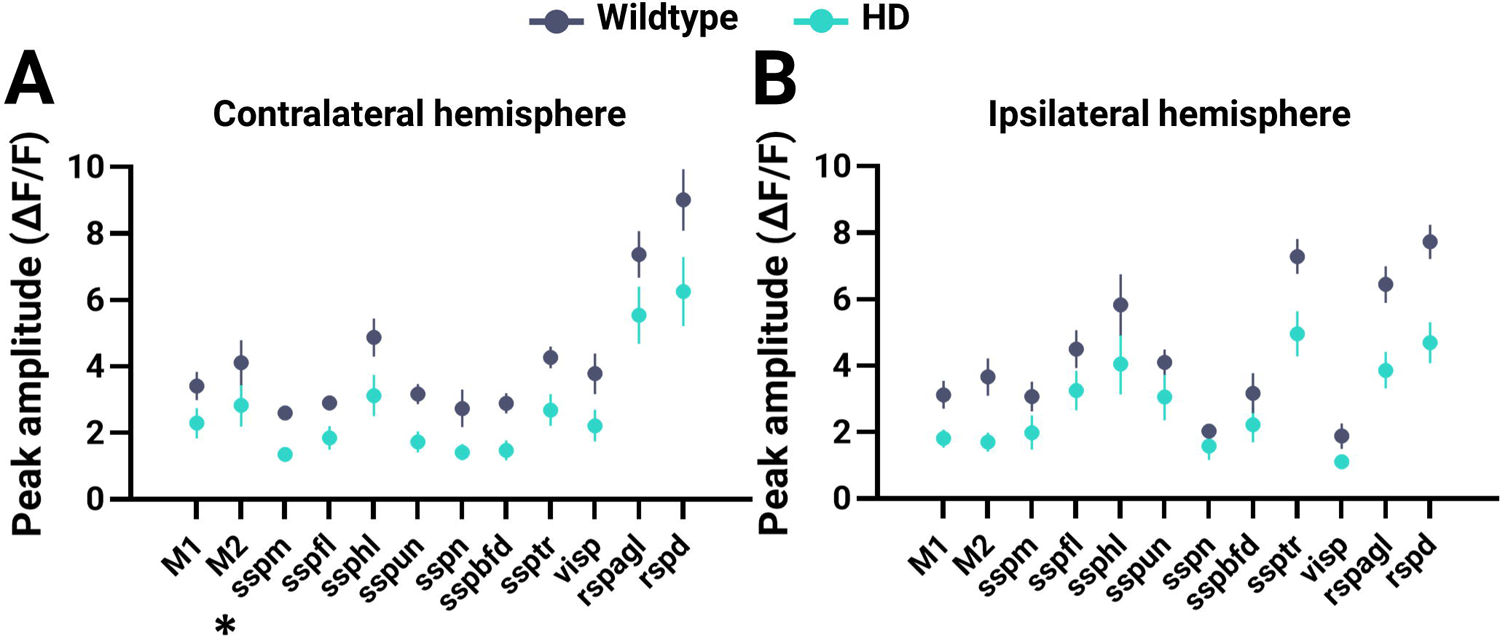
Mesoscale GCaMP imaging of the cortex during successful trials on Day 8. HD (n = 6) and WT (n = 4) mice are denoted in teal and gray, respectively. **(A-B)** Peak amplitude of regions of interest on Day 8 for successful trials in the contralateral (F_1,8_=5.925, p=0.0409, ANOVA)**(A)** and ipsilateral (F_1,8_=5.967, p=0.0404, ANOVA)**(B)** hemisphere. * denotes p=0.0290.

**Extended data Figure 5-2:**
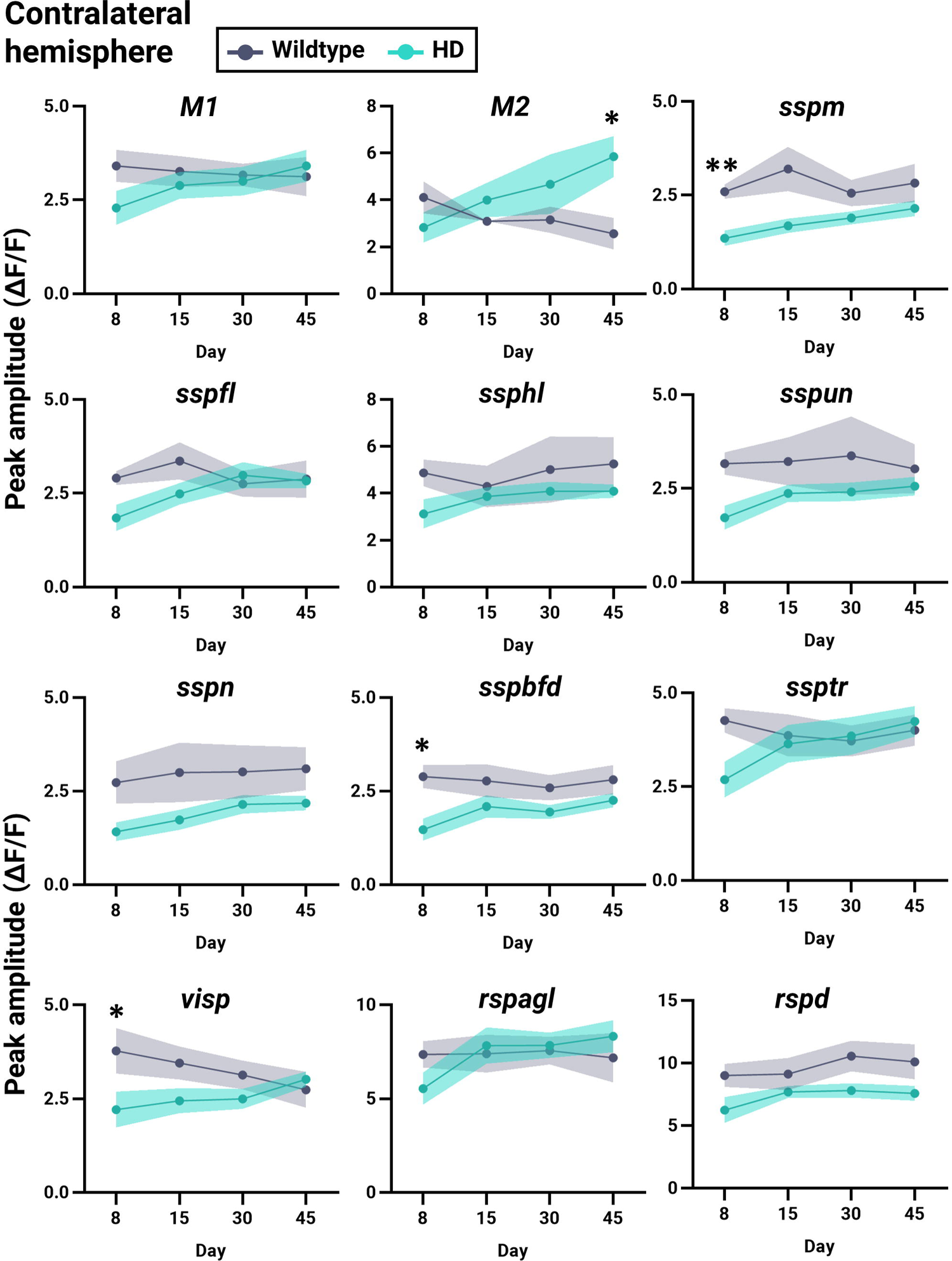
Change in contralateral hemisphere ROI peak ΔF/F amplitude over time. Peak ΔF/F amplitude of ROIs in the contralateral hemisphere over time for WT (n = 4)(gray) and HD (n = 6)(teal) mice. ** and * denotes p<0.01 and <0.05, respectively.

**Extended data Figure 5-3:**
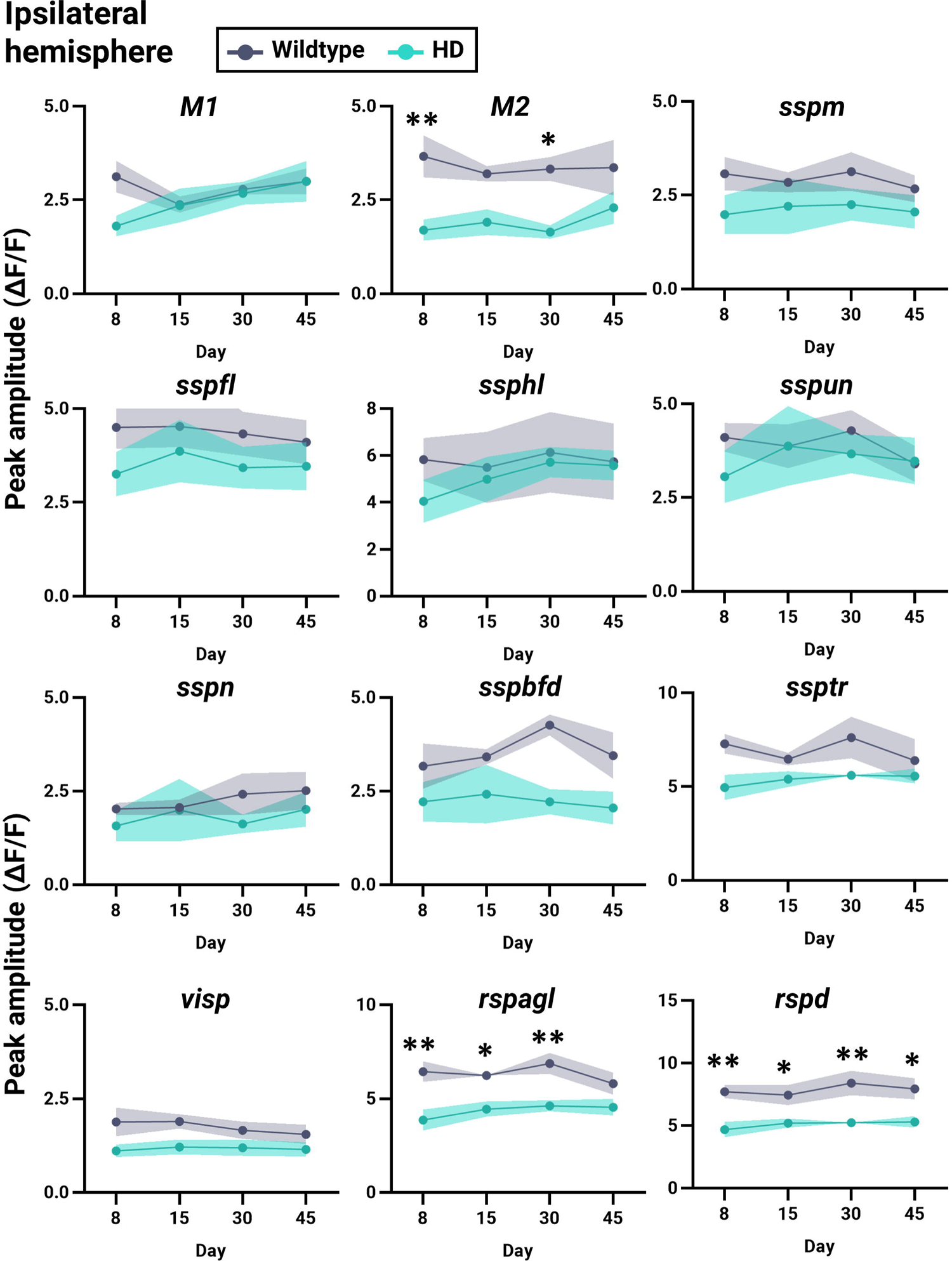
Change in ipsilateral hemisphere ROI peak ΔF/F amplitude over time. Peak ΔF/F amplitude of ROIs in the ipsilateral hemisphere over time for WT (n = 4)(gray) and HD (n = 6)(teal) mice. ** and * denotes p<0.01 and <0.05, respectively.

**Extended data Figure 6-1:**
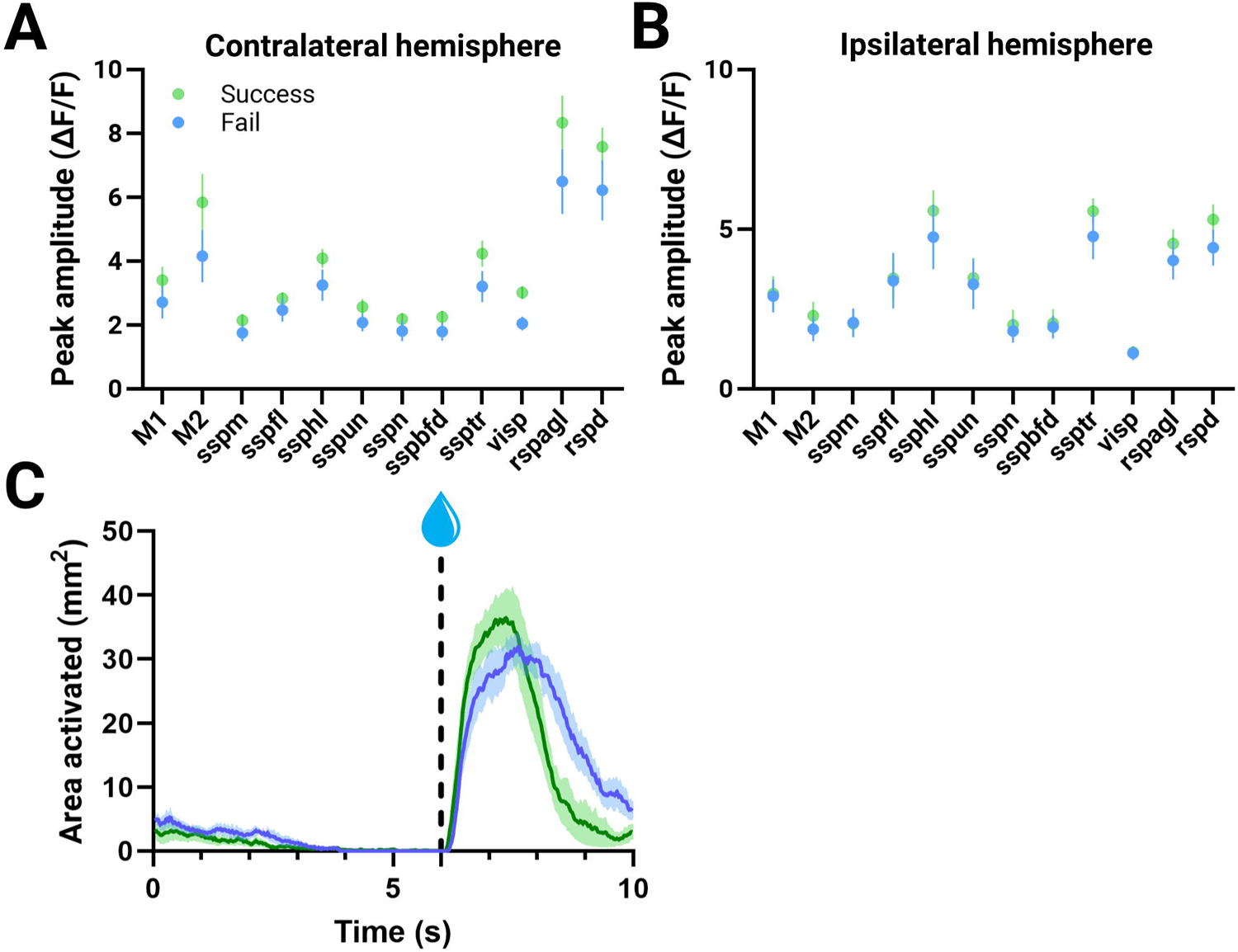
Mesoscale GCaMP imaging of the cortex during success and fail trials performed by HD mice on Day 45. HD (n = 6) success and fail trials are denoted in green and blue, respectively. **(A-B)** Peak amplitude of regions of interest in the contralateral (F_1,10_=2.540, p=0.1421, ANOVA)**(A)** and ipsilateral (F_1,10_=0.257, p=0.6230, ANOVA)**(B)** hemisphere for different trial types. **(C)** Area activated across the entire trial duration for successful and failed trials. The threshold was set at 4x standard deviation (STD) of the baseline. No significance between trial types.

## Acknowledgements

This work was supported by resources made available through the NeuroImaging and NeuroComputation Centre at the Djavad Mowafaghian Centre for Brain Health (RRID: SCR_019086); the UBC Vice-President Research and Innovation funding for the Dynamic Brain Circuits Cluster of Excellence; Canadian Institutes of Health Research grants FDN 143210 to L.A.R. and FDN 143209 to T.H.M and Brain Canada Vectrology Foundry to L.A.R. and T.H.M. T.H.M. is also supported by the Brain Canada Neurophotonics Platform, the Heart and Stroke Foundation of Canada, and the Fondation LeDucq. Y.W. is supported by the Vanier Canada Graduate Scholarship and UBC’s Four Year Doctoral Fellowship. We thank Pumin Wang for surgical assistance and Lily Zhang for technical genotyping assistance. We thank Evan Fung for Day 23 behavioral trial scoring. We thank Daniel Ramandi for expert advice on rotarod assessment and water-reaching behavioral camera setup. We further thank Daniel Ramandi and Jeffrey M. LeDue for expert advice on custom Matlab code writing for GCaMP analysis.

## Author contributions

Y.W. and L.A.R. designed the behavior and post-mortem experiment testing scheme. D.X. and T.H.M. designed the trial structure for the water-reaching task. Y.W. performed the research. Y.W. and M.D.S. analyzed the data. Y.W. wrote the manuscript with input from T.H.M., L.A.R., M.D.S. and D.X..

## Conflict of interest

The authors declare no competing financial interests.

## Notes

### Competing Interest Statement

The authors have declared no competing interest.

## References

1. Abada Y-SK, Schreiber R, Ellenbroek B (2013) Motor, emotional and cognitive deficits in adult BACHD mice: a model for Huntington’s disease. Behav Brain Res 238:243–251.

2. Albin RL, Reiner A, Anderson KD, Dure LS 4th, Handelin B, Balfour R, Whetsell WO Jr, Penney JB, Young AB (1992) Preferential loss of striato-external pallidal projection neurons in presymptomatic Huntington’s disease. Ann Neurol 31:425–430.

3. Ardesch DJ, Balbi M, Murphy TH (2017) Automated touch sensing in the mouse tapered beam test using Raspberry Pi. J Neurosci Methods 291:221–226.

4. Arnoux I et al. (2018) Metformin reverses early cortical network dysfunction and behavior changes in Huntington’s disease. eLife 7 Available at: http://dx.doi.org/10.7554/elife.38744.

5. Barry J, Akopian G, Cepeda C, Levine MS (2018) Striatal Direct and Indirect Pathway Output Structures Are Differentially Altered in Mouse Models of Huntington’s Disease. J Neurosci 38:4678–4694.

6. Batka RJ, Brown TJ, Mcmillan KP, Meadows RM, Jones KJ, Haulcomb MM (2014) The need for speed in rodent locomotion analyses. Anat Rec 297:1839–1864.

7. Becker MI, Calame DJ, Wrobel J, Person AL (2020) Online control of reach accuracy in mice. J Neurophysiol 124:1637–1655.

8. Brooks S, Higgs G, Janghra N, Jones L, Dunnett SB (2012) Longitudinal analysis of the behavioural phenotype in YAC128 (C57BL/6J) Huntington’s disease transgenic mice. Brain Res Bull 88:113–120.

9. Brunner C, Grillet M, Sans-Dublanc A, Farrow K, Lambert T, Macé E, Montaldo G, Urban A (2020) A Platform for Brain-wide Volumetric Functional Ultrasound Imaging and Analysis of Circuit Dynamics in Awake Mice. Neuron 108:861–875.e7 Available at: http://dx.doi.org/10.1016/j.neuron.2020.09.020.

10. Burgold J, Schulz-Trieglaff EK, Voelkl K, Gutiérrez-Ángel S, Bader JM, Hosp F, Mann M, Arzberger T, Klein R, Liebscher S, Dudanova I (2019) Cortical circuit alterations precede motor impairments in Huntington’s disease mice. Scientific Reports 9 Available at: http://dx.doi.org/10.1038/s41598-019-43024-w.

11. Cepeda C, Levine MS (2022) Synaptic dysfunction in Huntington’s disease: Lessons from genetic animal models. Neuroscientist 28:20–40.

12. Churchland MM, Afshar A, Shenoy KV (2006) A central source of movement variability. Neuron 52:1085–1096.

13. Cummings DM, André VM, Uzgil BO, Gee SM, Fisher YE, Cepeda C, Levine MS (2009) Alterations in cortical excitation and inhibition in genetic mouse models of Huntington’s disease. J Neurosci 29:10371–10386.

14. Czajkowski R, Jayaprakash B, Wiltgen B, Rogerson T, Guzman-Karlsson MC, Barth AL, Trachtenberg JT, Silva AJ (2014) Encoding and storage of spatial information in the retrosplenial cortex. Proc Natl Acad Sci U S A 111:8661–8666.

15. Dana H, Chen T-W, Hu A, Shields BC, Guo C, Looger LL, Kim DS, Svoboda K (2014) Thy1-GCaMP6 transgenic mice for neuronal population imaging in vivo. PLoS One 9:e108697.

16. Dhawale AK, Miyamoto YR, Smith MA, Ölveczky BP (2019) Adaptive Regulation of Motor Variability. Curr Biol 29:3551–3562.e7.

17. Donzis EJ, Estrada-Sánchez AM, Indersmitten T, Oikonomou K, Tran CH, Wang C, Latifi S, Golshani P, Cepeda C, Levine MS (2020) Cortical Network Dynamics Is Altered in Mouse Models of Huntington’s Disease. Cereb Cortex 30:2372–2388.

18. Elliott D, Helsen WF, Chua R (2001) A century later: Woodworth’s (1899) two-component model of goal-directed aiming. Psychol Bull 127:342–357.

19. Feigin A, Ghilardi M-F, Huang C, Ma Y, Carbon M, Guttman M, Paulsen JS, Ghez CP, Eidelberg D (2006) Preclinical Huntington’s disease: compensatory brain responses during learning. Ann Neurol 59:53–59.

20. Fernández-García S, Conde-Berriozabal S, García-García E, Gort-Paniello C, Bernal-Casas D, García-Díaz Barriga G, López-Gil J, Muñoz-Moreno E, Soria G, Campa L, Artigas F, Rodríguez MJ, Alberch J, Masana M (2020) M2 cortex-dorsolateral striatum stimulation reverses motor symptoms and synaptic deficits in Huntington’s disease. Elife 9 Available at: http://dx.doi.org/10.7554/eLife.57017.

21. Galiñanes GL, Bonardi C, Huber D (2018) Directional Reaching for Water as a Cortex-Dependent Behavioral Framework for Mice. Cell Rep 22:2767–2783.

22. Guo J-Z, Sauerbrei BA, Cohen JD, Mischiati M, Graves AR, Pisanello F, Branson KM, Hantman AW (2021) Disrupting cortico-cerebellar communication impairs dexterity. Elife 10 Available at: http://dx.doi.org/10.7554/eLife.65906.

23. Heming EA, Cross KP, Takei T, Cook DJ, Scott SH (2019) Independent representations of ipsilateral and contralateral limbs in primary motor cortex. Elife 8 Available at: http://dx.doi.org/10.7554/eLife.48190.

24. Klein A, Sacrey L-AR, Dunnett SB, Whishaw IQ, Nikkhah G (2011) Proximal movements compensate for distal forelimb movement impairments in a reach-to-eat task in Huntington’s disease: New insights into motor impairments in a real-world skill. Neurobiology of Disease 41:560–569 Available at: http://dx.doi.org/10.1016/j.nbd.2010.11.002.

25. Koch ET, Sepers MD, Cheng J, Raymond LA (2022) Early Changes in Striatal Activity and Motor Kinematics in a Huntington’s Disease Mouse Model. Mov Disord Available at: http://dx.doi.org/10.1002/mds.29168.

26. Liu H, Zhang C, Xu J, Jin J, Cheng L, Miao X, Wu Q, Wei Z, Liu P, Lu H, van Zijl PCM, Ross CA, Hua J, Duan W (2021) Huntingtin silencing delays onset and slows progression of Huntington’s disease: a biomarker study. Brain 144:3101–3113 Available at: http://dx.doi.org/10.1093/brain/awab190.

27. MacDonald ME et al. (1993) A novel gene containing a trinucleotide repeat that is expanded and unstable on Huntington’s disease chromosomes. Cell 72:971–983.

28. Mathis A, Mamidanna P, Cury KM, Abe T, Murthy VN, Mathis MW, Bethge M (2018) DeepLabCut: markerless pose estimation of user-defined body parts with deep learning. Nat Neurosci 21:1281–1289.

29. McColgan P, Tabrizi SJ (2018) Huntington’s disease: a clinical review. Eur J Neurol 25:24–34.

30. McFadyen MP, Kusek G, Bolivar VJ, Flaherty L (2003) Differences among eight inbred strains of mice in motor ability and motor learning on a rotorod. Genes Brain Behav 2:214–219.

31. Menalled L, Lutz C, Ramboz S, Brunner D, Lager B, Noble S, Park L, Howland D (2014) A field guide to working with mouse models of Huntington’s Disease. Bar Harbor: PsychoGenics Inc, The Jackson Laboratory, CHDI Foundation.

32. Milczarek MM, Vann SD, Sengpiel F (2018) Spatial Memory Engram in the Mouse Retrosplenial Cortex. Curr Biol 28:1975–1980.e6.

33. Mohammed H, Li Y, Di Grazia P, Bernstein A, Agger S, Hollis E (2020) Temporal regulation of motor behavior on a modified forelimb dexterity test in mice. bioRxiv:2020.10.18.344507 Available at: https://www.biorxiv.org/content/10.1101/2020.10.18.344507 [Accessed July 24, 2022].

34. Murphy TH, Boyd JD, Bolaños F, Vanni MP, Silasi G, Haupt D, LeDue JM (2016) High-throughput automated home-cage mesoscopic functional imaging of mouse cortex. Nat Commun 7:11611.

35. Pallier PN, Drew CJG, Jennifer Morton A (2009) The detection and measurement of locomotor deficits in a transgenic mouse model of Huntington’s disease are task- and protocol-dependent: Influence of non-motor factors on locomotor function. Brain Research Bulletin 78:347–355 Available at: http://dx.doi.org/10.1016/j.brainresbull.2008.10.007.

36. Peng Q, Wu B, Jiang M, Jin J, Hou Z, Zheng J, Zhang J, Duan W (2016) Characterization of Behavioral, Neuropathological, Brain Metabolic and Key Molecular Changes in zQ175 Knock-In Mouse Model of Huntington’s Disease. PLoS One 11:e0148839.

37. Pinto L, Rajan K, DePasquale B, Thiberge SY, Tank DW, Brody CD (2019) Task-Dependent Changes in the Large-Scale Dynamics and Necessity of Cortical Regions. Neuron 104:810– 824.e9.

38. Pouladi MA, Morton AJ, Hayden MR (2013) Choosing an animal model for the study of Huntington’s disease. Nat Rev Neurosci 14:708–721.

39. Quarta E, Scaglione A, Lucchesi J, Sacconi L, Allegra Mascaro AL, Pavone FS (2022) Distributed and Localized Dynamics Emerge in the Mouse Neocortex during Reach-to-Grasp Behavior. J Neurosci 42:777–788.

40. Reiner A, Albin RL, Anderson KD, D’Amato CJ, Penney JB, Young AB (1988) Differential loss of striatal projection neurons in Huntington disease. Proc Natl Acad Sci U S A 85:5733– 5737.

41. Sauerbrei BA, Guo J-Z, Cohen JD, Mischiati M, Guo W, Kabra M, Verma N, Mensh B, Branson K, Hantman AW (2020) Cortical pattern generation during dexterous movement is input-driven. Nature 577:386–391.

42. Sepers MD, Mackay JP, Koch E, Xiao D, Mohajerani MH, Chan AW, Smith-Dijak AI, Ramandi D, Murphy TH, Raymond LA (2021) Altered cortical processing of sensory input in Huntington disease mouse models. bioRxiv:2021.07.18.452688 Available at: https://www.biorxiv.org/content/10.1101/2021.07.18.452688 [Accessed March 19, 2022].

43. Shabbott B, Ravindran R, Schumacher JW, Wasserman PB, Marder KS, Mazzoni P (2013) Learning fast accurate movements requires intact frontostriatal circuits. Front Hum Neurosci 7:752.

44. Silasi G, Xiao D, Vanni MP, Chen ACN, Murphy TH (2016) Intact skull chronic windows for mesoscopic wide-field imaging in awake mice. J Neurosci Methods 267:141–149.

45. Smith, Rocha, McLean (2014) Progressive axonal transport and synaptic protein changes correlate with behavioral and neuropathological abnormalities in the heterozygous Q175 KI mouse model …. Hum Mol Genet Available at: https://academic.oup.com/hmg/article-abstract/23/17/4510/736560.

46. Sofroniew NJ, Flickinger D, King J, Svoboda K (2016) A large field of view two-photon mesoscope with subcellular resolution for in vivo imaging. Elife 5 Available at: http://dx.doi.org/10.7554/eLife.14472.

47. Soma S, Yoshida J, Kato S, Takahashi Y, Nonomura S, Sugimura YK, Ríos A, Kawabata M, Kobayashi K, Kato F, Sakai Y, Isomura Y (2019) Ipsilateral-Dominant Control of Limb Movements in Rodent Posterior Parietal Cortex. J Neurosci 39:485–502.

48. Southwell AL, Smith-Dijak A, Kay C, Sepers M, Villanueva EB, Parsons MP, Xie Y, Anderson L, Felczak B, Waltl S, Ko S, Cheung D, Dal Cengio L, Slama R, Petoukhov E, Raymond LA, Hayden MR (2016) An enhanced Q175 knock-in mouse model of Huntington disease with higher mutant huntingtin levels and accelerated disease phenotypes. Hum Mol Genet 25:3654–3675.

49. Steinmetz NA, Zatka-Haas P, Carandini M, Harris KD (2019) Distributed coding of choice, action and engagement across the mouse brain. Nature 576:266–273.

50. Vanni MP, Chan AW, Balbi M, Silasi G, Murphy TH (2017) Mesoscale Mapping of Mouse Cortex Reveals Frequency-Dependent Cycling between Distinct Macroscale Functional Modules. J Neurosci 37:7513–7533.

51. Vanni MP, Murphy TH (2014) Mesoscale transcranial spontaneous activity mapping in GCaMP3 transgenic mice reveals extensive reciprocal connections between areas of somatomotor cortex. J Neurosci 34:15931–15946.

52. Wekselblatt JB, Flister ED, Piscopo DM, Niell CM (2016) Large-scale imaging of cortical dynamics during sensory perception and behavior. J Neurophysiol 115:2852–2866.

53. Whishaw IQ, Agha BM, Kuntz JR, Qandeel, Faraji J, Mohajerani MH (2018a) Tongue protrusions modify the syntax of skilled reaching for food by the mouse: Evidence for flexibility in action selection and shared hand/mouth central modulation of action. Behavioural Brain Research 341:37–44 Available at: http://dx.doi.org/10.1016/j.bbr.2017.12.006.

54. Whishaw IQ, Faraji J, Agha BM, Kuntz JR, Metz GAS, Mohajerani MH (2018b) A mouse’s spontaneous eating repertoire aids performance on laboratory skilled reaching tasks: A motoric example of instinctual drift with an ethological description of the withdraw movements in freely-moving and head-fixed mice. Behavioural Brain Research 337:80–90 Available at: http://dx.doi.org/10.1016/j.bbr.2017.09.044.

55. Whishaw IQ, Pellis SM, Gorny BP (1992) Skilled reaching in rats and humans: evidence for parallel development or homology. Behav Brain Res 47:59–70.

56. Woodard CL, Bolaños F, Boyd JD, Silasi G, Murphy TH, Raymond LA (2017) An Automated Home-Cage System to Assess Learning and Performance of a Skilled Motor Task in a Mouse Model of Huntington’s Disease. eneuro 4:ENEURO.0141–17.2017 Available at: http://dx.doi.org/10.1523/eneuro.0141-17.2017.

57. Woodard CL, Sepers MD, Raymond LA (2021) Impaired Refinement of Kinematic Variability in Huntington Disease Mice on an Automated Home Cage Forelimb Motor Task. J Neurosci 41:8589–8602.

58. Xiao D, Forys BJ, Vanni MP, Murphy TH (2021) MesoNet allows automated scaling and segmentation of mouse mesoscale cortical maps using machine learning. Nat Commun 12:5992.

59. Xiao D, Vanni MP, Mitelut CC, Chan AW, LeDue JM, Xie Y, Chen AC, Swindale NV, Murphy TH (2017) Mapping cortical mesoscopic networks of single spiking cortical or sub-cortical neurons. Elife 6 Available at: http://dx.doi.org/10.7554/eLife.19976.

